# Genetic introgression is a catalyst for diversification in an Old World fruit bat radiation

**DOI:** 10.64898/2026.02.10.702973

**Authors:** Balaji Chattopadhyay, Kritika M. Garg, Susan M. Tsang, Joanna L. Coleman, Reizl P. Jose, Faisal Ali Bin Anwarali Khan, Sigit Wiantoro, Ian H. Mendenhall, Gavin J.D. Smith, Frank E. Rheindt

## Abstract

Genetic introgression, the exchange of genes between closely related species, was once thought to be rare and counterproductive to species differentiation until more recent evidence showed that introgression is pervasive in nature and may even facilitate the transfer of important functional traits among species. However, evidence for the role of introgression in facilitating speciation in natural systems is rare. In this study, we investigated the role of introgression in accelerating trait sharing and speciation in a poorly understood group of South and Southeast Asian fruit bats of the genus *Cynopterus* using genome-wide DNA from 93 individuals, including both fresh and historic museum specimens. We uncovered substantial new species level diversity within the genus and retrieved multiple strong signatures of complex genetic introgression, including two putative events of hybrid speciation within this radiation. In the genomic areas implicated in most of these introgression events, we identified a locus potentially linked to sperm integrity and fusion of egg and sperm. Further, we observed high between-species divergence in an area of the genome linked to skull development. The *Cynopterus* radiation augments a small but growing list of examples showing that genetic introgression – rather than forming an impediment against differentiation – is likely to boost and reinforce diversification processes, for example through the transfer of functional loci that may facilitate innovations. Our study also highlights the importance of museum collections in understanding cryptic diversity and speciation in complex biogeographic settings.

---

Our planet’s organismal diversity is the result of evolutionary processes that have produced lineages across a spectrum spanning from species-poor to extremely species-rich. The mechanisms underlying this diversification and its unequal outcomes in different parts of the tree-of-life have long been a matter of biological inquiry. Over the past ∼100 years, geographic and reproductive isolation have been regarded as the main prerequisites for successful speciation: Ernst Mayr’s biological species concept, rooted in the allopatric speciation paradigm (Mayr 1942), emphasized the prohibitive effects that even small amounts of gene flow may have on the divergence of an ancestral species into two daughter species. More recently, examples of sympatric speciation have been identified in which an ancestral lineage can diverge even if the two daughter lineages remain in geographic overlap with each other (Barluenga et al. 2006), but most of these scenarios are restricted to unusual spatial circumstances or lifestyles (e.g. host-dependent parasites (Nosil 2007), etc.).

With the advent of genome-wide DNA datasets, it has become increasingly obvious that gene flow often remains a widespread though rare occurrence even after two species have diverged, and frequently results in the genetic introgression of alleles from one species into the other (Green et al. 2010; Rheindt and Edwards 2011; Harrison and Larson 2014; Chattopadhyay et al. 2016; Taylor and Larson 2019). Genomic datasets have identified genetic introgression in many species in which it had never been suspected (e.g. hominids (Green et al. 2010; Medugorac et al. 2017; Racimo et al. 2015; Rocha et al. 2023)).

Even so, there is still substantial uncertainty about the role that genetic introgression may play in the differentiation process. In the Mayrian speciation framework, genetic introgression is a rare by-product of evolution that occasionally counteracts the differentiation process. However, recent studies have shown that introgression events could lead to the infiltration of alleles that may confer new traits or functionalities in populations (Huerta-Sánchez et al. 2014; Norris et al. 2015; Garg et al. 2019; Rossi et al. 2024). Examples of this phenomenon remain scarce, but – if widespread – it may suggest that genetic introgression may be an active driver of speciation in some organisms. In *Heliconius* butterflies, for instance, differences in wing color among populations are mediated by introgression from sympatric populations of other species for the purpose of Müllerian mimicry (Pardo-Diaz et al. 2012), promoting differentiation within species. Other species-rich radiations, including the classic Darwin’s finches (*Geospiza* spp. (Grant and Grant 2008)), are also known for their high incidence of introgression, suggesting that the latter is unlikely to have slowed differentiation in this group.

In this study, we used fresh and historic DNA samples, the latter sourced from museum specimens up to 113 years old, to sequence datasets of genome-wide DNA and investigate the impact of genetic introgression on the diversification of a poorly-known radiation of tropical Asian fruit bats of the genus *Cynopterus*. Our phylogenomic and population-genomic inferences show the need for a complete taxonomic revision of a group whose species diversity had been grossly underestimated by traditional methods. More importantly, we document a history of rampant introgressive gene flow among *Cynopterus* bats across their entire range, including two possible hybrid speciation events in which daughter species display a mosaic of genomic contributions from two putative parent species. We identify specific genes in elevated “introgression hotspots” with an important role in reproduction. Our results suggest that genetic introgression is not merely a rare side-product of *Cynopterus* diversification but has actively driven the speciation process and produced an assemblage that may have been much poorer if species boundaries were less porous.

## MATERIALS AND METHODS

### Sampling and DNA extraction

We used a combination of freshly collected and historic museum samples to ensure range-wide representation of the *Cynopterus brachyotis* complex (n = 88) and closely related species (Table S1). Five outgroup taxa were also included in our genome-wide data generation (Table S1). We obtained fresh tissue samples from either wing biopsies or muscle (Table S1). For historic samples, we used variable sources of tissue such as muscle, dried tissue from skulls (crusties), and dried wing samples (Table S1). We used the DNeasy blood and tissue kit (Qiagen, Germany), following the manufacturer’s protocol, to extract high molecular weight DNA from fresh samples. All historic samples were extracted in a dedicated ancient DNA facility away from fresh samples. Each sample was handled with fresh gloves and instruments to avoid contamination. For historic samples, we used a modified DNeasy blood and tissue extraction protocol with double the volumes of buffer and digestion times (Chattopadhyay et al. 2019). We utilized MinElute columns (Qiagen, Germany) to capture small single-stranded DNA fragments. As historic DNA is heavily fragmented, the use of columns for DNA purification substantially improved the yield of extracted DNA. For all extractions we included dedicated negative controls, and final extracts were quantified using high sensitivity Qubit kits (Thermo Fischer Scientific, USA). DNA from historic samples was further run on an AATI Fragment Analyser (Agilent, USA) to obtain a distribution of fragment sizes.

### Target enrichment marker design

A novel approach was used to design target loci, such that both phylogenomic and population genomic information could be obtained from the same set of markers. Details of the target locus design have been provided by Chattopadhyay et al. (2019). In brief, we identified 1,165 single copy conserved exons shared between the *Cynopterus brachyotis* (GCA_009793145.1; Chattopadhyay et al. 2020) and *Myotis lucifugus* genomes (GCA_000147115.1). We used EvolMarkers (Li et al. 2012) to identify these exons, which had to be at least 200 bp in length. For each exon identified in *C. brachyotis*, we further isolated the two adjoining stretches of DNA sequence up to 500 bp upstream and downstream of the exon. We checked for overlapping regions and merged them in bedtools 2.28.0 (Quinlan and Hall 2010). The target loci were further filtered for GC content, such that only loci with an average GC content between 30 to 60% were retained. We checked for repeat regions using RepeatMasker 4.0.7 (Smit et al. 2015) and removed loci with long tracts of repeat regions. Any locus consisting of less than 0.2% repeat elements was selected and soft-masked. In addition, we included 19 introns (Teeling et al. 2005) and five sperm protein genes that are important for mammalian reproduction (Swanson and Vacquier 2002) to shed light on *Cynopterus* evolution. This locus selection regime resulted in 1,184 loci comprising approximately 1.5 Mbp of sequence data, or slightly less than 0.1% of the *Cynopterus brachyotis* genome. We designed RNA probes for hybrid capture using Mycoarray (USA). In brief, we designed 100 bp probes with a 4X tilling density to ensure overlap between probe sequences in order to maximize the length of target loci retrieved from degraded museum samples.

### Library preparation and hybrid capture

For fresh samples, we used the NEB Ultra II kit (New England BioLabs, USA) for library preparation following the manufacturer’s protocol. Prior to library preparation, the samples were sheared using a Bioruptor pico (Diagenode, Belgium). A total of 100μl of the sample was subjected to 13 cycles of sonication, with each cycle consisting of 30s of sonication followed by 30s of rest. This sonication regime yielded fragments of ∼ 250bp. For barcoding the libraries, we used dual 8bp barcodes. For library preparation of historic DNA, we followed Chattopadhyay et al. (2019). In brief, samples were first treated with the FFPE DNA repair kit (New England BioLabs, USA) to reduce DNA damage commonly observed in historic samples, and then prepared using the NEB Ultra II kit. We used negative controls for all library preparations. The final libraries were quantified using a Qubit and an AATI Fragment Analyzer.

We enriched for the target loci using the hybrid capture protocol of Chattopadhyay et al. (2019). In brief, we used a modification of the myBaits (Arbor Biosciences, USA) version 3 protocol for historic samples whereby diluted baits were used at 50% strength and hybridization was carried out at 60^0^ C for 40 hours, with separate hybridization reactions for each historic sample. For fresh samples, we pooled three sample libraries at an equimolar concentration and carried out hybridization at 65^0^ C for 20 hours. After hybridization, samples were cleaned following the myBaits protocol and PCR enrichment was performed for clean enriched libraries using IS5 and IS6 primers (Kircher et al. 2012). The enriched libraries were cleaned using Ampure beads (Beckman Coulter Life Sciences, USA) and quantified using both a Qubit and an AATI Fragment Analyzer. Finally, enriched libraries were pooled at an equimolar concentration and run on the Illumina 4000 platform. We carried out 150bp paired-end runs, with separate lanes for fresh and ancient samples.

### NGS data cleanup and data matrix

We obtained a total of ∼1.2 billion raw reads. We used FastQC 0.11.7 (Andrews 2010) to check for sequence quality and adapter contamination. Reads with bases below a PHRED score of 10 had been removed by the service provider. Both cutadapt 1.12 (Martin 2011) and Trimmomatic 0.38 (Bolger et al. 2014) were used to remove low quality reads and adapter contamination. We then again performed a quality check in FastQC to ensure the complete removal of adapters. Following this, PCR duplicates were removed using the dedupe program within bbmap 36.84 (Bushnell 2014). We performed an additional cleanup for historic samples, as they are prone to DNA damage, by identifying damage patterns and rescaling the bam file quality scores using mapDamage 2 (Jónsson et al. 2013). The rescaled bam files were converted to fastq files and 10bp from both the 5 prime and 3 prime ends were removed using Seqtk 1.2-r94 (https://github.com/lh3/seqtk). These cleaned reads were used for all further analyses.

The cleaned reads formed the basis for analyses based both on DNA sequences and SNPs. For sequence-based analysis, the target loci were assembled using the HybPiper 1.2 pipeline with default settings (Johnson et al. 2018). Within this pipeline, the cleaned reads were mapped to target loci using the Burrows-Wheeler Aligner 0.7.12 (Li and Durbin 2009) and then assembled using SPAdes 3.11.1 (Prjibelski et al. 2020). We provided exon-intron boundaries to obtain complete target locus sequences. We used the scripts get_seq_lengths.py and hybpiper_stats.py within the pipeline to assess missing data levels and target locus length for the assembled data. Finally, the retrieve_sequence.py script was used to obtain sequences for each locus across samples in fasta format. We obtained sequence data for 1,182 out to 1,184 target loci.

For SNP-based analysis, we generated two different datasets (SNP dataset I using the *de novo* genome of *C. brachyotis*, GCA_009793145.1, and SNP dataset II using a pseudo-chromosomal assembly of *C. brachyotis* obtained from Chromosomer). Chromosomer generates pseudo-chromosomal assemblies by aligning scaffolds to a closely related chromosomal assembly. In our case the closest genome assembly available at the time was that of *Rhinolophus ferrumequinum* (GCF_004115285.1; Jebb et al. 2020). We were able to map 24,399 scaffolds (806,702,397 bp) to the *R. ferrumequinum* genome, resulting in a partial pseudo-chromosomal assembly of *Cynopterus brachyotis*. To avoid potential artifacts from degraded DNA, we only included fresh *Cynopterus* samples along with the outgroup *Ptenochirus jagori* for SNP calling, as all deep lineages happened to be represented by fresh samples. For both datasets, the reads were mapped using the Burrows-Wheeler Aligner and then sorted using SAMtools 1.9 (Li et al. 2009; Li 2011). We used ANGSD 0.923-3-ga8ed56f [24] to identify SNPs using a p-value cut-off of 1E-6, with a minimum read depth of 10, PHREAD score ≥ 30, and less than 20% missing data. We then filtered SNPs to remove linked loci using the indep-pairwise algorithm within PLINK 1.9 (Purcell et al. 2007), with a sliding window size of 50 SNPs, a step size of 10 and an *r*^2^ correlation coefficient cut-off of 0.9. We also removed any loci not in Hardy–Weinberg equilibrium using PLINK while correcting the p-value for multiple comparisons. Finally, we removed any loci under selection using the R 3.5.3 (R Core Team 2018) package pcadapt (Luu et al. 2017) with a 10% false recovery rate.

### Phylogenomic analyses

For phylogenomic analysis, we included sequence data from eight publicly available bat genomes (*Eonycteris spelaea*, GCA_003508835.1, Wen et al. (2018); *Pteropus vampyrus*, GCA_000151845.2; *Pteropus alecto*, GCA_000325575.1, Zhang et al. (2013); *Eidolon helvum*, GCA_000465285.1; *Rhinolophus sinicus*, GCA_001888835.1, Dong et al. (2014); *Megaderma lyra*, GCA_004026885.1, Parker et al. (2013); *Hipposideros armiger*, GCA_001890085.1, Dong et al. (2014); and *Craseonycteris thonglongyai*, GCA_004027555.1). Target loci were isolated from these genomes using a script provided by Chattopadhyay et al. (2020) (https://github.com/gargkritika/append_sequences_to_existing_alignments). Sequences were aligned using auto settings in MAFFT v7.130b (Katoh and Standley 2016), allowing reverse complementation whenever required. Ten historic samples were removed from our dataset due to poor data quality. Unfortunately, this resulted in the loss of data for two *Cynopterus* taxa (*C. titthaecheilus* and *C. horsfieldii princeps*). The remaining 91 samples including outgroups were used for phylogenetic analysis. The aligned data were cleaned using Gblocks 0.91b (Castresana 2020), allowing for a maximum of 50% missing data for any conserved and flanking position, and default settings for other parameters. The average locus length was around 1.2 kb (range: 160 to 7,817bp). Analyses were not only performed on the ‘most-inclusive dataset’ as outlined above, but also on a dataset containing only fresh samples to account for any bias due to the degradation of DNA observed in museum samples.

We used both classical concatenation and coalescent-based species tree methods (Edwards et al. 2007) to reconstruct the phylogeny of *Cynopterus* species. For concatenation analysis, loci were concatenated using the AMAS pipeline (Borowiec 2016), and a maximum likelihood tree was generated using the rapid bootstrap method in RAxML 8.2 (Stamatakis 2014), with 1,000 bootstrap runs under the GTR+GAMMA model for sequence evolution. For species tree reconstruction, we used MP-EST 1.6 (Liu et al. 2010) following Liu et al. (2017). Individual gene trees were estimated using RAxML within the phyluce pipeline with 100 bootstraps per gene tree and *Hipposideros armiger*, an insectivorous bat, as an outgroup. Rooting was performed online on the STRAW server (https://straw.phylolab.net/SpeciesTreeAnalysis/index.php). MP-EST analysis was based on 980 loci for which outgroup sequences were available. The gene trees were separated into 100 bootstrap files and species trees were estimated for each bootstrap file in MP-EST. The 100 bootstrap species trees were then used to estimate nodal support using the majority-rule consensus method in Phylip v 3.69 (Felsenstein 2005). Final trees of all phylogenomic analyses were visualized in FigTree v1.42 (http://tree.bio.ed.ac.uk/software/figtree/).

Even after multiple cleanup steps during the target enrichment process, the mitochondrial genome is often sequenced as ‘by-catch’ due to its high copy number within a cell. Using the CLC workbench 9.5 (https://www.qiagenbioinformatics.com/), we took advantage of this feature and assembled consensus sequences for all samples, each with a minimum read depth of 10, spanning the complete mitochondrial cyt *b* gene (1,140 bp). These sequences were aligned with cyt *b* sequences already available for the genus *Cynopterus* on Genbank (see Table S2 for accession IDs). We used the *Rousettus amplexicaudatus* sequence from the Philippines as an outgroup, and identified HKY+I+G as the best substitution model using jmodeltest 2.1.10 (Posada 2008; Darriba et al. 2012). We performed two runs in MrBayes v3.2.6 (Ronquist et al. 2012) with four chains each, using default settings for swapping frequency and temperature. The program was run for 10,000,000 generations while the standard deviation of the split frequency remained below 0.01. We checked for convergence in Tracer v1.6 (http://tree.bio.ed.ac.uk/software/tracer/) and constructed a consensus tree after a 25% burn-in.

Using FastaChar (Merckelbach and Borges 2020), we further identified private mutations for the cyt *b* gene among all the lineages of the genus *Cynopterus* identified in this study. A private mutation is defined as a mutation that is present in all individuals of a lineage of interest but not in other lineages. Individuals which were shown to be subject to mito-nuclear discordance were removed.

### Coalescent modeling

We used a coalescent approach to estimate population divergence times and migration rates in G-PhoCS (Gronau et al. 2011). We only considered fresh samples for this analysis, thereby avoiding any bias resulting from DNA damage of historic samples. We mapped the 1,182 loci to the *C. brachyotis* scaffold assembly and only considered loci at least 20kb apart. We then removed any loci under selection and recombination, resulting in a dataset of 482 loci (582,430 bp) for coalescent modeling. We carried out multiple trial runs in G-PhoCS considering various gene flow scenarios. For all runs we used the topology obtained from RAxML as a baseline. We ran three models in G-PhoCS to investigate the evolutionary trajectory surrounding two potential polytomies. For the first model, we accepted the tree topology based on RAxML (regardless of branch support) and allowed for gene flow between lineages. For the second and third models, we tested for evolutionary scenarios that are more consistent with true (‘hard’) polytomies at the relevant nodes (see supporting information). We performed two independent runs for each model, each for 1 million generations, and consistently checked for convergence using TRACER. When estimating evolutionary parameters, we performed a 50% burn-in and noted mean values as well as the 95% confidence range for each parameter. Migration was only inferred if the 95% confidence estimate of a parameter did not overlap with zero and the minimum value was greater than 0.01 in both runs.

### Introgression analysis

We employed three approaches to understand the role of genetic introgression in shaping the diversification of the genus *Cynopterus*. First, we calculated four-taxon ABBA-BABA statistics using the ANGSD package. ABBA-BABA is a well-established method to differentiate between introgression and incomplete lineage sorting. We only used fresh samples for this analysis and considered sites with a mapping quality and PHRED score ≥30. To test for significance, we performed jackknifing of 20-kb blocks. Significant secondary admixture was inferred when Z scores exceeded −3 or +3. We performed ABBA-BABA analysis for both SNP datasets.

Second, we used DSuite (Malinsky et al. 2021) to calculate D statistics with the MP-EST species tree as a guide. We computed Patterson’s D and Dmin statistics to understand patterns of gene flow. The Dmin is the minimum value of a D statistic possible by rearranging trios. This parameter is useful in cases where phylogenetic relationships between taxa are poorly defined. Again, we inferred significant D statistic values whenever Z scores exceeded 3. We further computed f-branch statistics to identify any introgression events between internal tree branches. We only used the first SNP dataset for this analysis.

Third, we used the admixture graph approach in qpbrute (Leathlobhair et al. 2018; Liu et al. 2019) to understand patterns of introgression. This approach is rooted in D-statistics and explores patterns of pairwise allele sharing to arrive at an ideal set of admixturegraphs inferring the relationships of target lineages along with streams of introgressive gene flow. We exclusively employed the second SNP dataset for this analysis because qpbrute can only handle chromosome-level or high quality genome assemblies with less than 100 scaffolds. Using the qpGraph program, part of ADMIXTOOLS 5.1 (Patterson et al. 2012), qpbrute employed a heuristic algorithm to iteratively fit complex models of admixture. From the models identified using qpGraph, qpbrute selected the best admixture graph using Bayes factor analysis.

### Identifying loci under introgression and important for speciation

To identify genomic regions with an elevated introgression signal, we employed a sliding window ABBA-BABA approach, using 100kb windows containing at least 50 SNPs per window. Using our first SNP dataset, we measured Patterson’s D, fdM (Malinsky et al. 2021), and fd (Martin et al. 2015) statistics using scripts provided by Simon Martin (https://github.com/simonhmartin/genomics_general). Among these, fdM is thought to be best suited to identifying windows of introgression, although it is known to be conservative. We homed in on any windows with an elevated introgression signal across 56 possible species combinations. For each combination, we selected the top 10% windows with an elevated signal of introgression and compared them across all combinations. We then selected elevated windows of introgression which were observed in at least 50% of the combinations and used bedtools to isolate genes present or partially present in them. Gene annotation was performed using a combination of Pannzer2 (Törönen et al. 2018), eggNogMapper v2 (Cantalapiedra et al. 2021), and the web version of InterProScan (Quevillon et al. 2005).

We also estimated pairwise Dxy values for all species pairs to identify regions of the genome which are highly differentiated. We used scripts provided by Simon Martin (https://github.com/simonhmartin/genomics_general) for the analysis identifying windows with the top 5% Dxy signal. Akin to introgression analysis, we selected elevated Dxy windows which were present in at least 50% of comparisons (n = 14) and short-listed the genes potentially involved in speciation. For all identified genes, we performed gene ontology (GO) enrichment analysis using ShinyGO 0.76 (Ge et al. 2020).

### Ancestry painting

For two putative hybrid species identified in our dataset (*C. minutus* and an undescribed lineage from Palawan, see Results) we performed ancestry painting to shed light on patterns of inheritance. We first identified the differentially fixed sites in the putative parent species and then compared the alleles in the putative hybrid species to the parents. We allowed for 10% missing data. The scripts used for this analysis were obtained from the following source: https://github.com/mmatschiner/tutorials/tree/master/analysis_of_introgression_with_snp_dat a. We isolated genes located in the vicinity of fixed loci using a window size of 2,000bp. We further performed GO enrichment analysis using ShinyGO 0.76 for these genes.

## RESULTS

### NGS data matrix

We obtained a total of ∼1.2 billion raw reads and retained ∼664 million reads after cleanup steps, with an average of 7 million paired-end reads (∼2.1 Gbp) per sample (Table S1). Our analyses were based on two categories of genome-wide DNA data: sequence loci (spanning approximately 1.2 kb on average) and single nucleotide polymorphisms (SNPs). The sequence data matrix obtained spanned 1,454,131 bp and was approximately 92% complete. To preclude ancient DNA bias, our SNP-based analyses exclusively encompassed fresh samples (56 ingroup individuals plus one outgroup). We obtained 331,534 and 172,049 SNPs for SNP dataset I and SNP dataset II respectively. After removing linked SNPs and those that deviated from Hardy-Weinberg equilibrium and neutrality, we retained 217,352 SNPs for the first SNP dataset and 107,836 SNPs for the second dataset (see Table S3 for details). The overall genotyping rate for the clean SNP data was ∼94 %.

### Phylogenomic reconstruction

We performed separate phylogenetic analysis for the mitochondrial cytochrome *b* (cyt *b*) gene and nuclear target loci. Our results, based on both nuclear and mitochondrial data, indicate a split of taxa traditionally placed within *C. brachyotis* into six deeply diverged and potentially species-level lineages, two of which (*C. minutus, C. ceylonensis*) are not even embedded within the remainder of the *C. brachyotis* complex (Fig. 1). The results from genome-wide DNA markers are largely consistent with previous suggestions based on short mitochondrial sequences (Campbell et al. 2004) (Fig. 1), and point to two new and likely undescribed lineages of *Cynopterus* from Palawan and Sulawesi respectively (Figs 1, S1, and S2). Based on cyt *b* data alone, two species-level clades of *C. horsfieldii*, one comprising the nominate subspecies from Java to Sumbawa and the other including populations from elsewhere for which the name *C. princeps* Miller, 1906, has priority, emerged in a non-sister position (Fig. 1, S1 and S2). The revised taxonomic treatment of the various taxa of the genus *Cynopterus* is summarized in Table 1.

**Figure 1:**
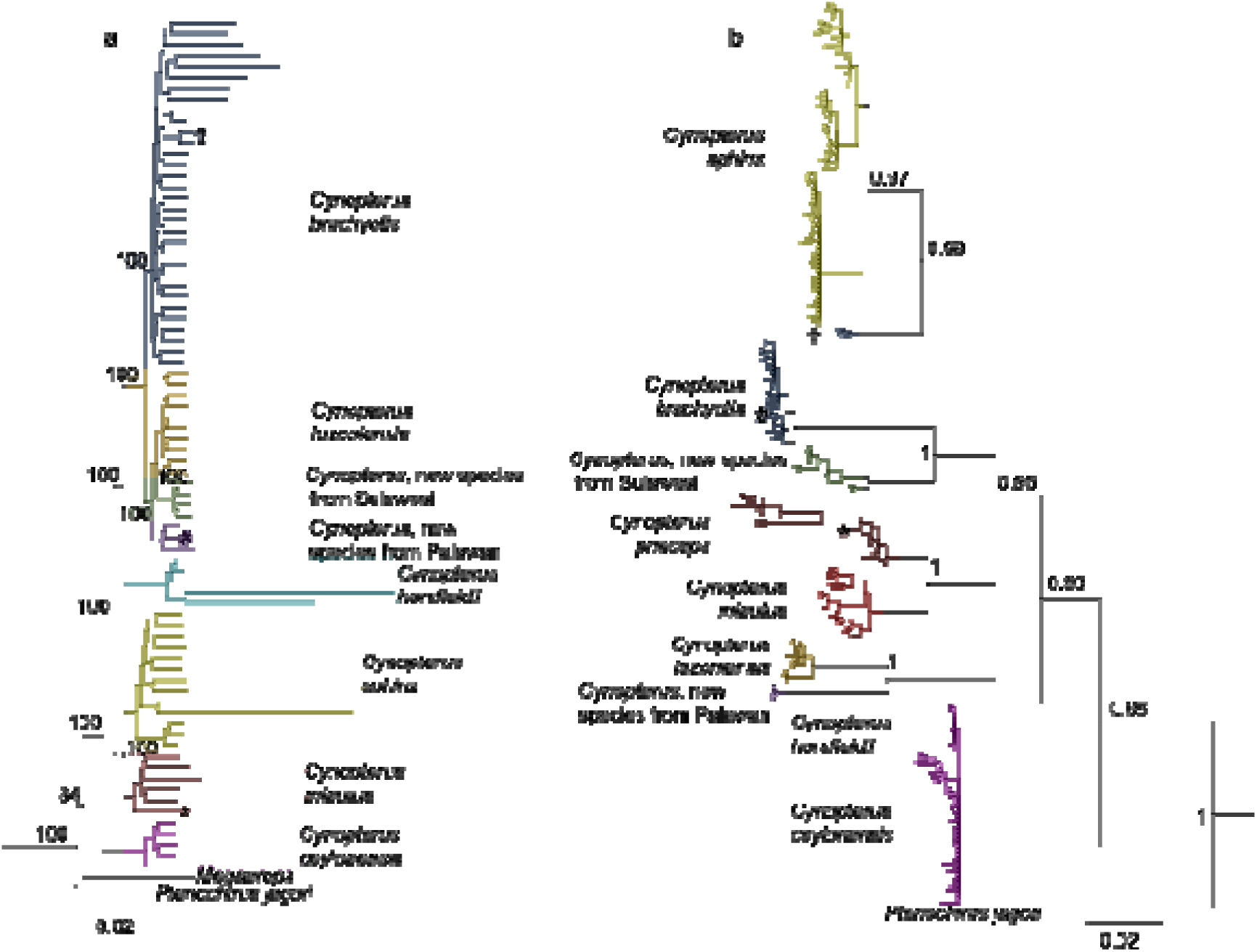
Phylogenomic relationships among members of the genus *Cynopterus*. a) Maximum likelihood tree based on concatenated nuclear loci (1,182 sequence loci; total alignment length 1,454,131 bp) generated using RAxML. Only outgroups *Megaerops* and *Ptenochirus jagori* shown for clarity; b) Bayesian tree based on 1,140 bp of the cytochrome *b* gene using MrBayes; only outgroup *Ptenochirus jagori* shown for clarity. Nodal values indicate bootstrap support (1a) and posterior probabilities (1b). Support values shown only for major nodes. Three major instances of mito-nuclear discordance are indicated with an asterisk, hash, and dagger symbol, respectively. The asterisk symbol denotes an individual with a *C. princeps* mitochondrial haplotype identified as *C. minutus* based on nuclear loci; the hash symbol denotes an individual with a *C. brachyotis* mitochondrial haplotype identified as the new *Cynopterus* lineage from Palawan based on nuclear loci; the dagger symbol denotes individuals with mitochondrial haplotypes close to *C. sphinx* but identified as *C. brachyotis* based on nuclear loci.

**Table 1:**
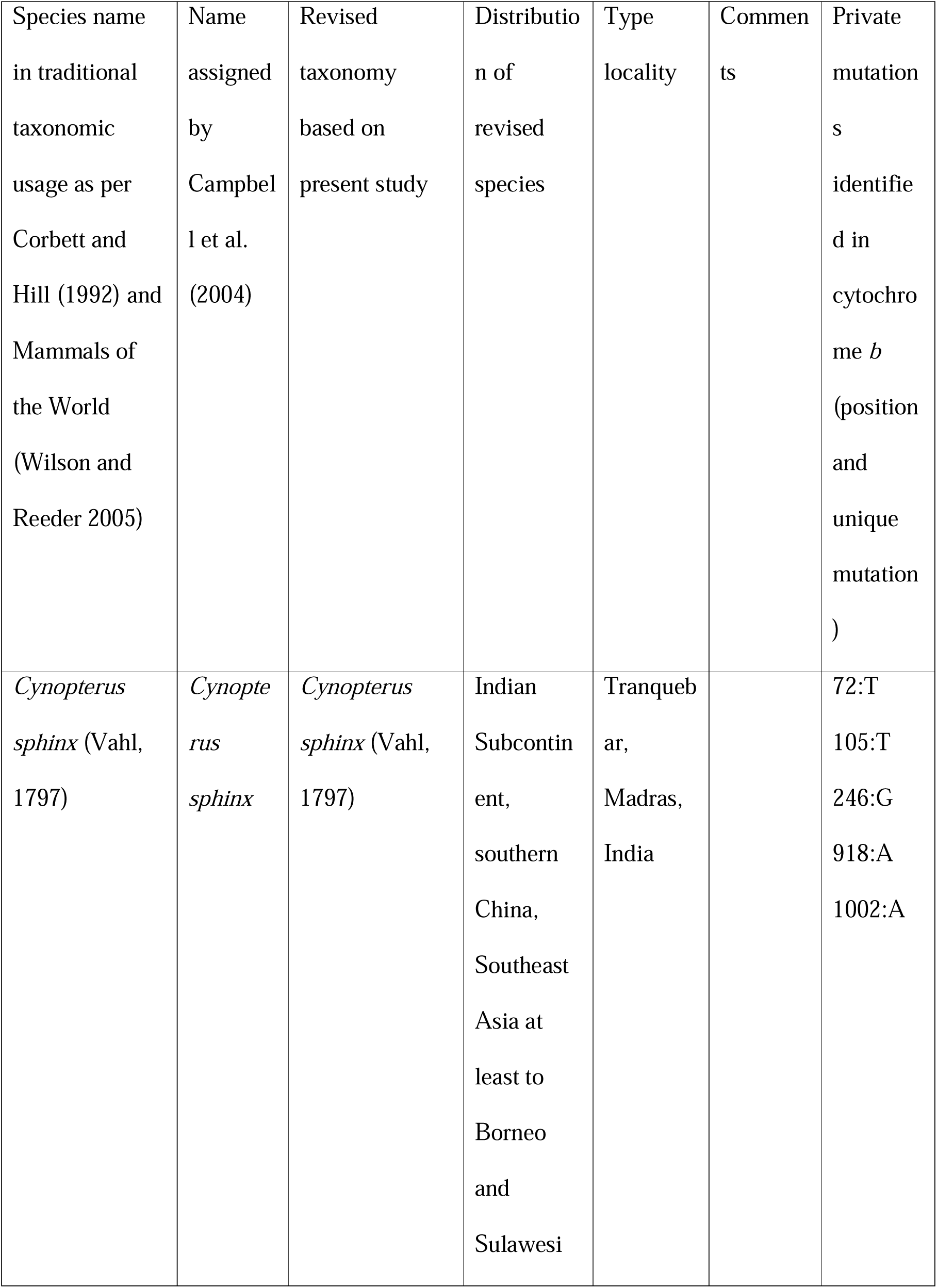
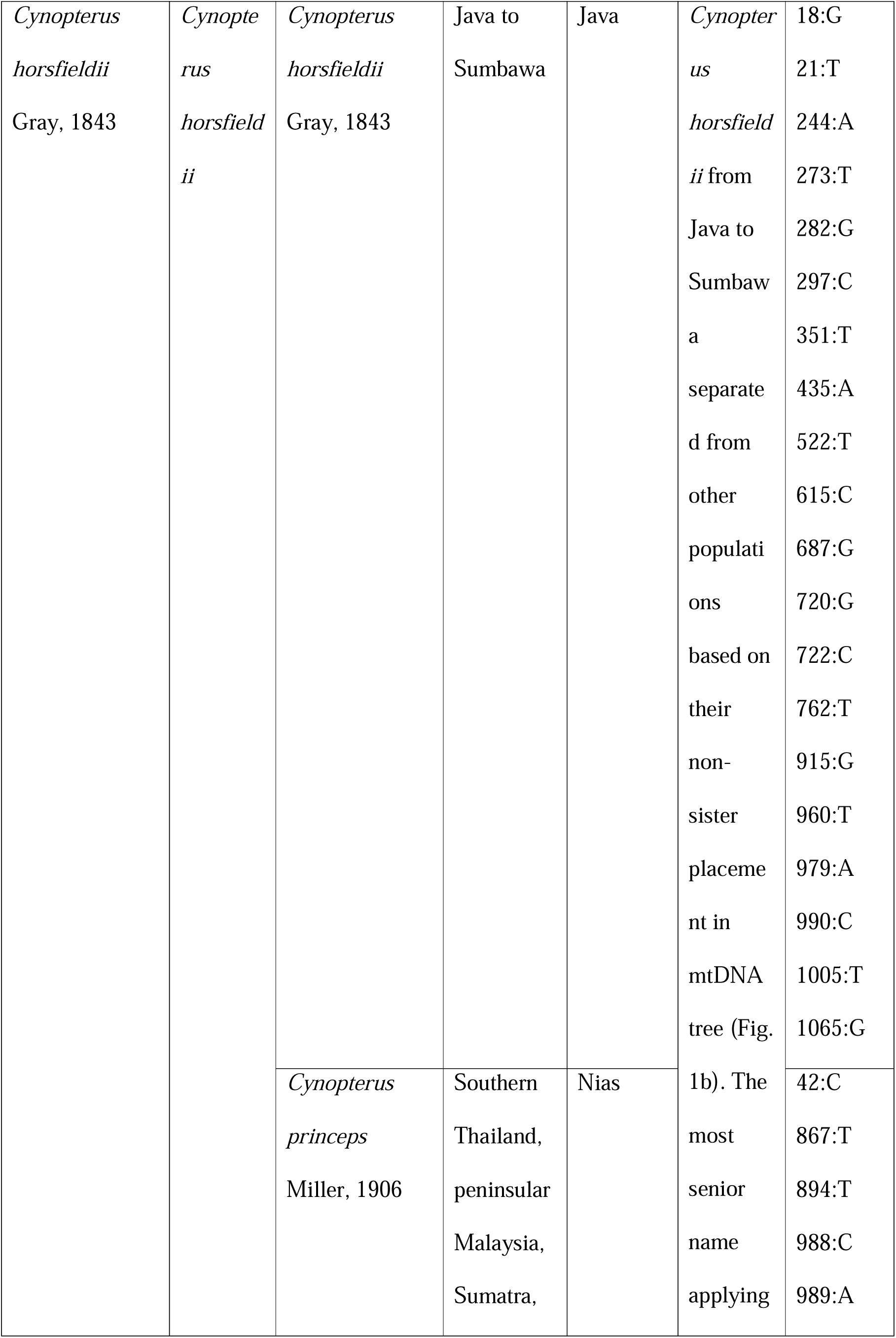
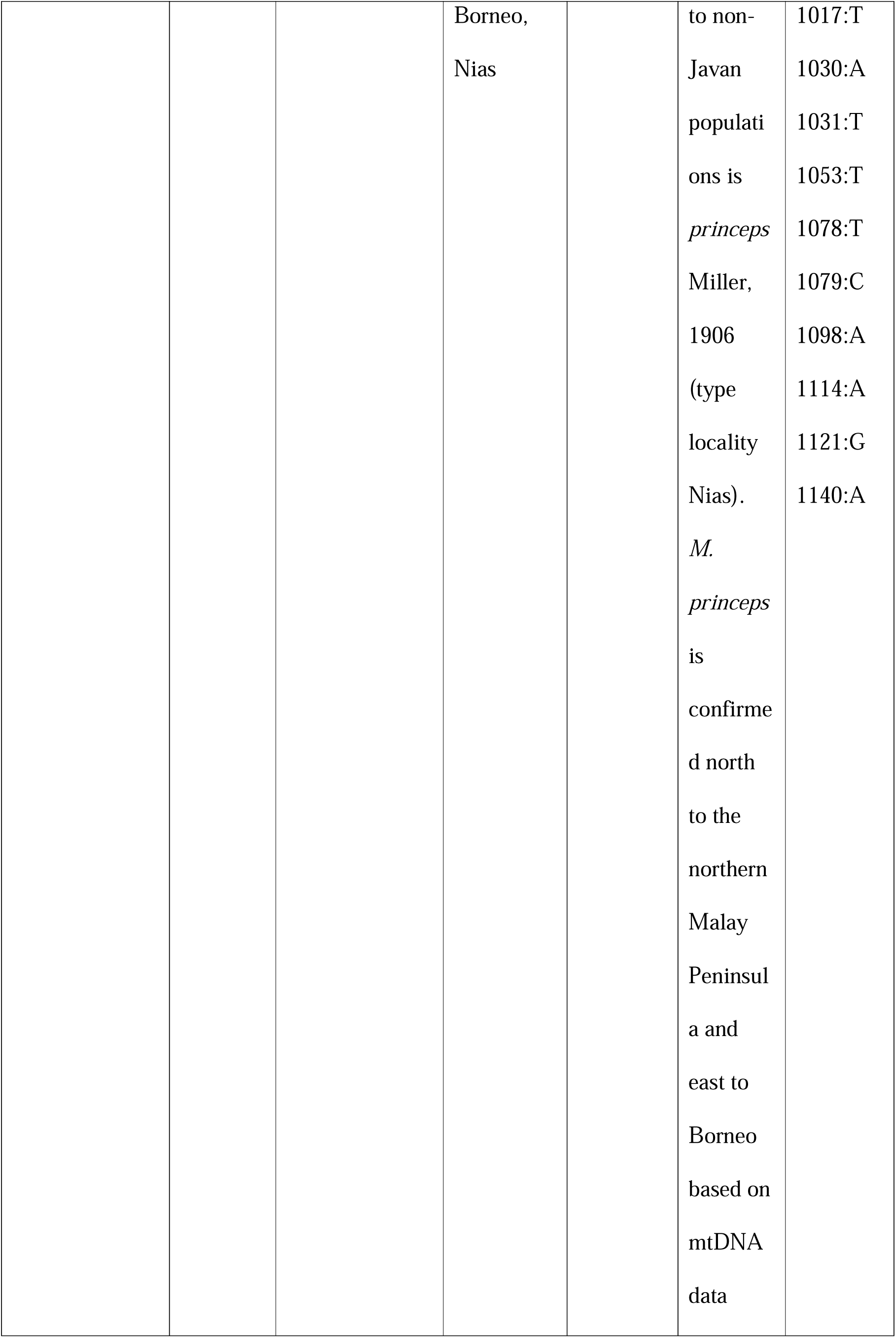
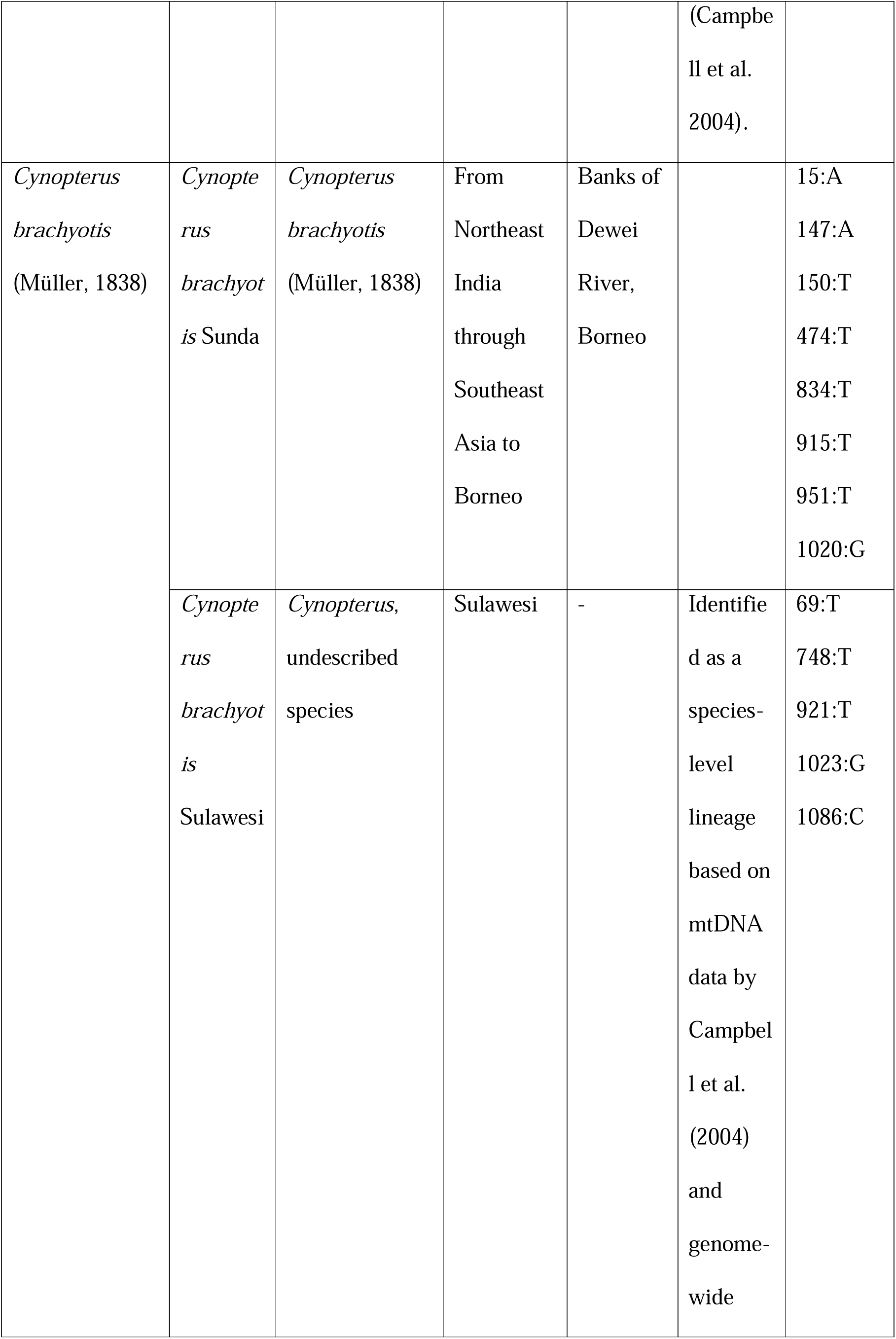
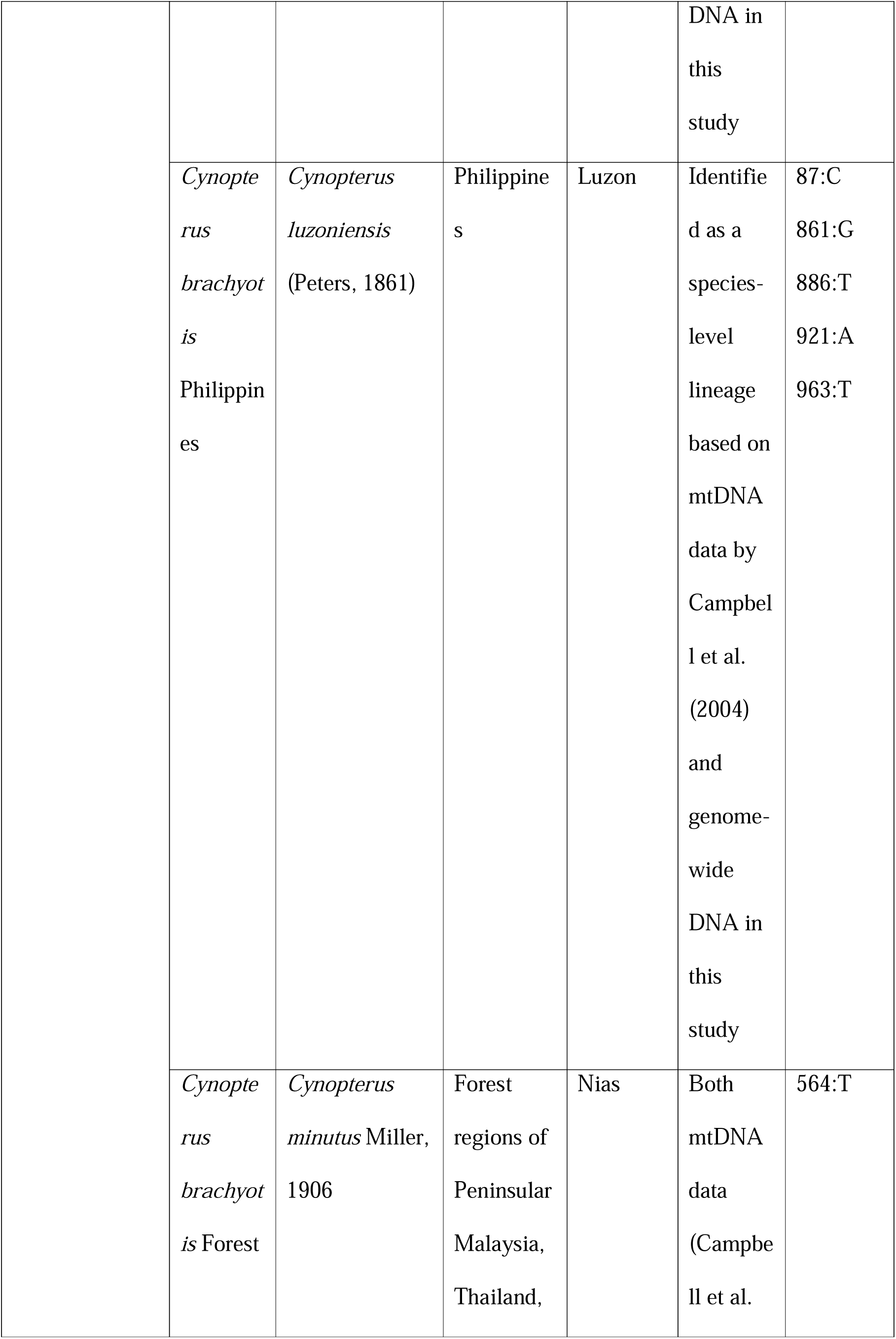
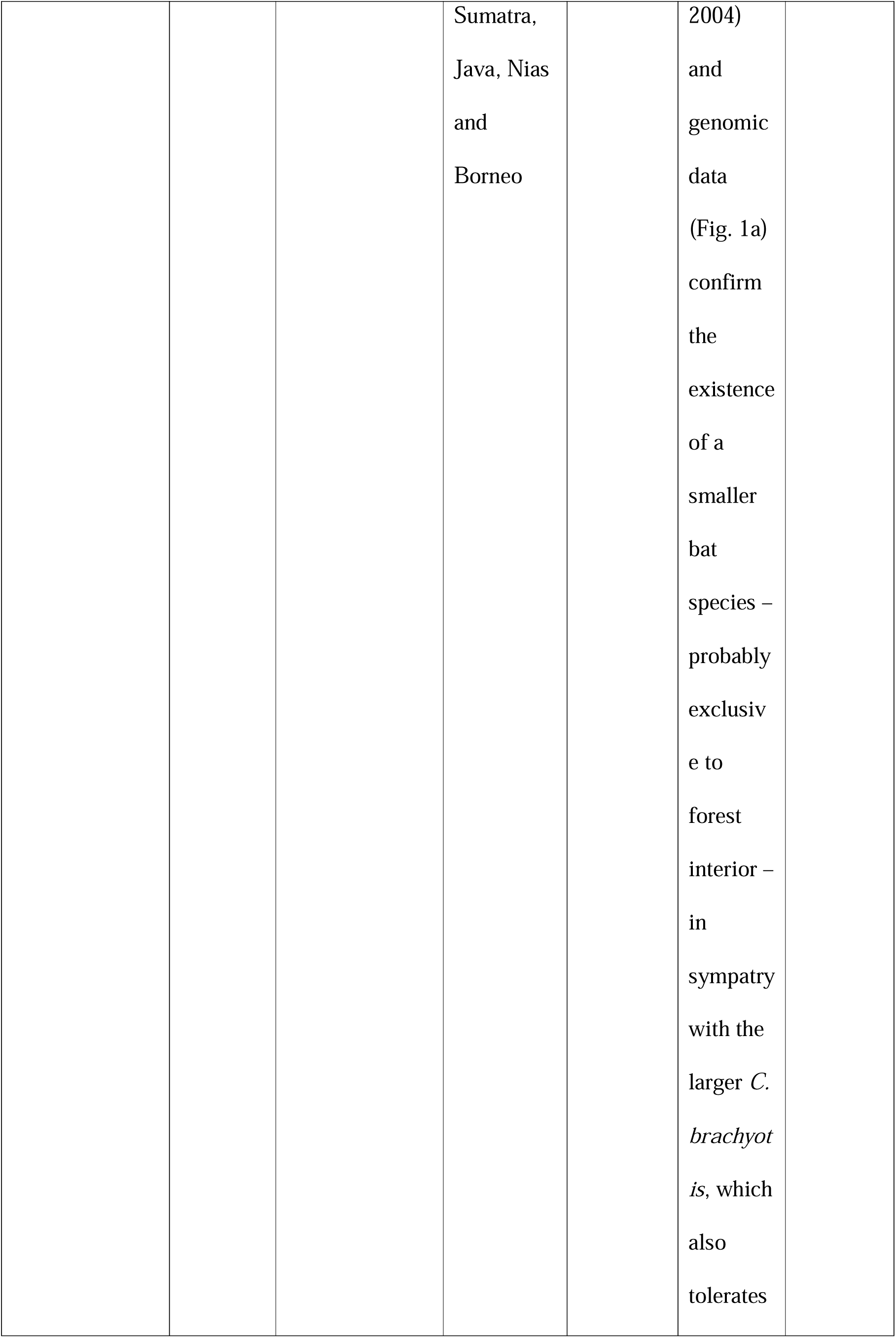
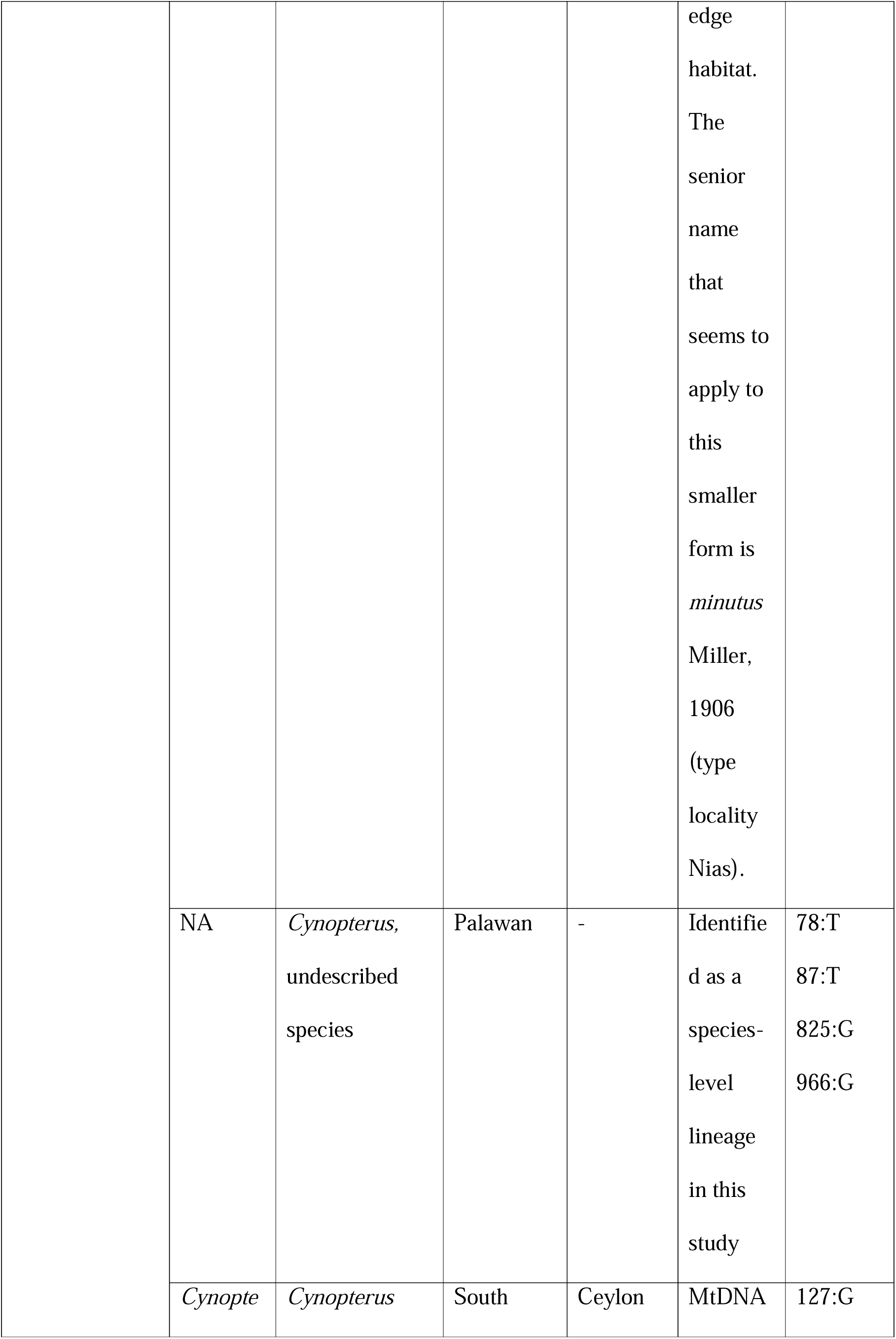
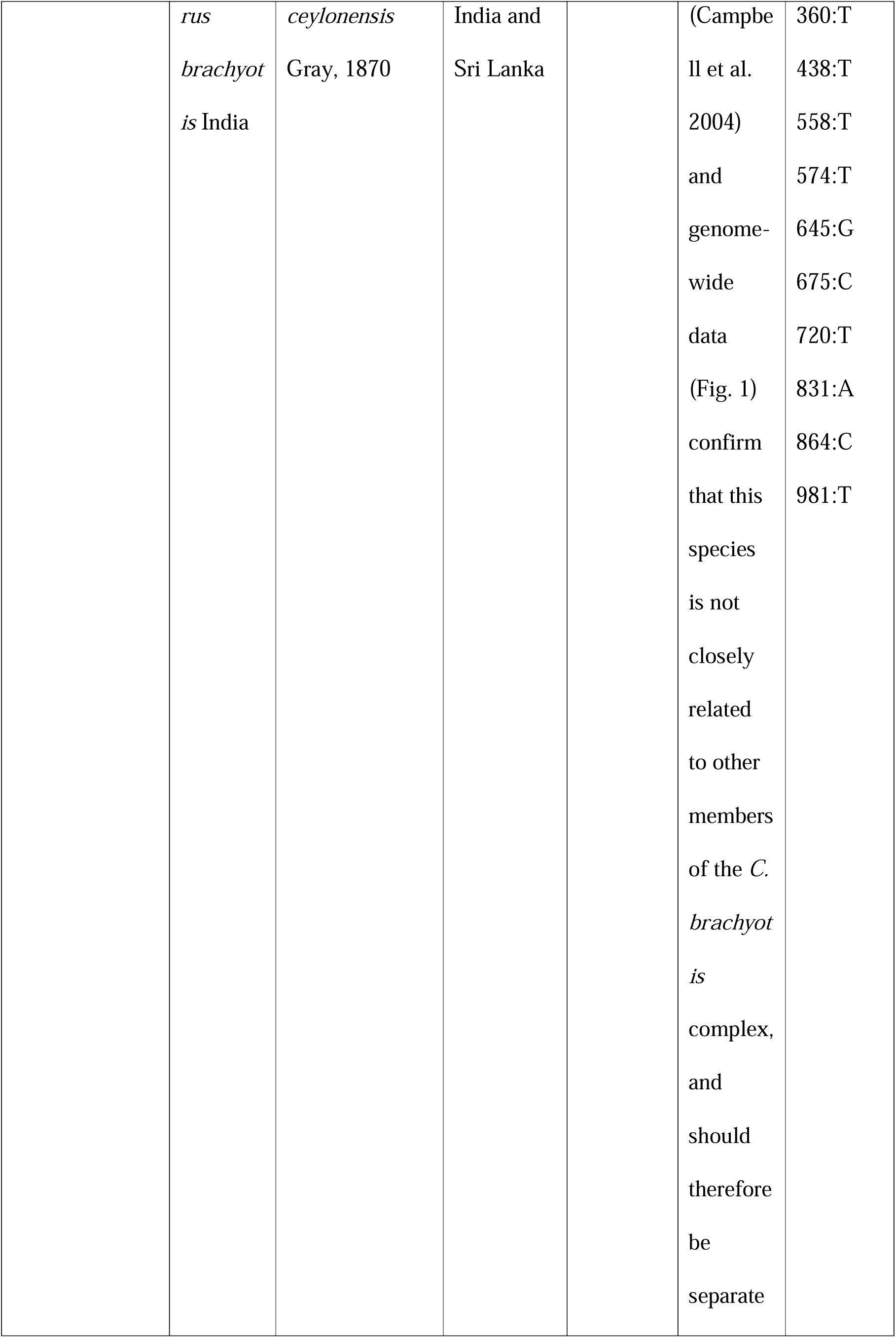
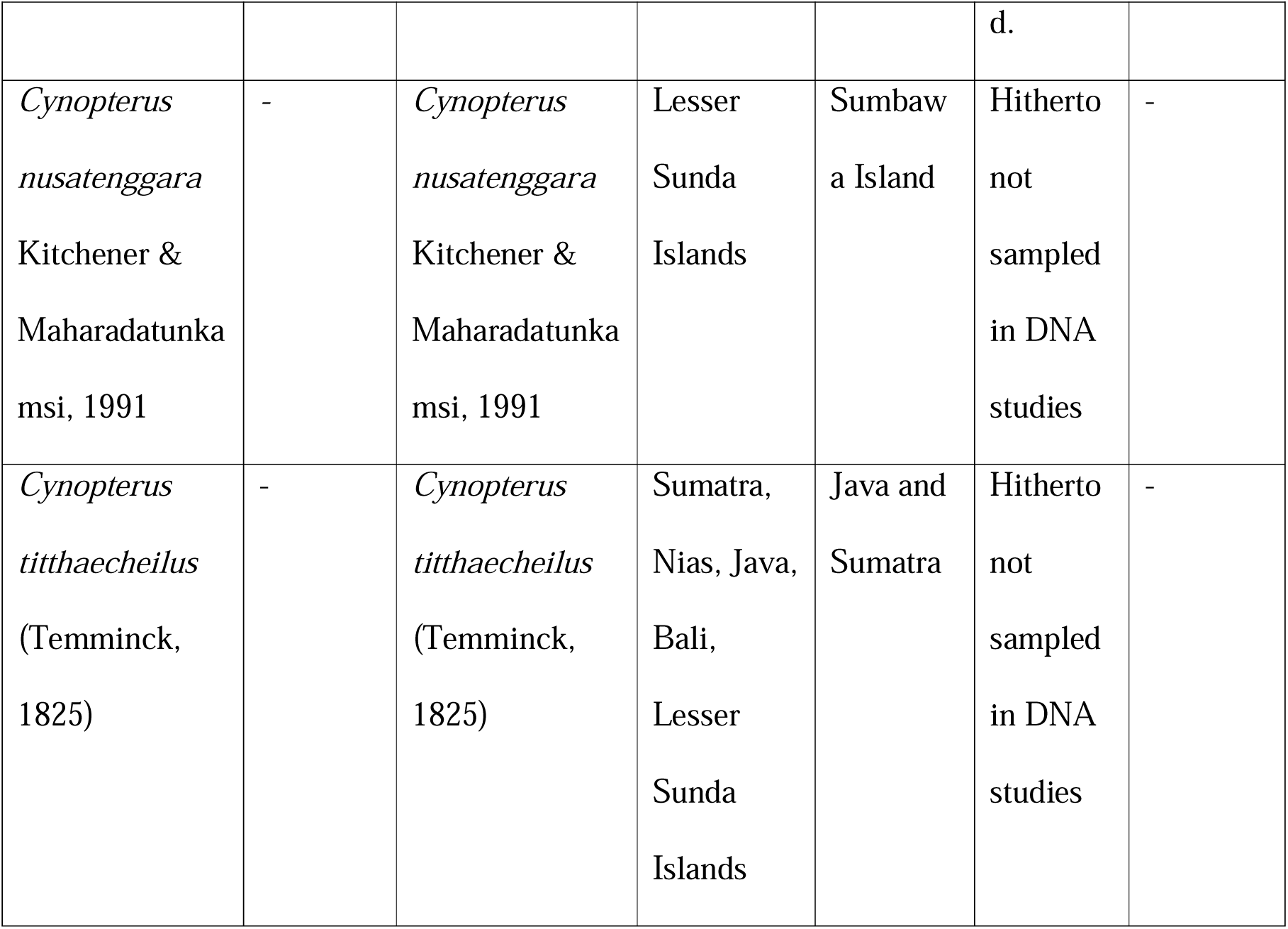
Revised taxonomic treatment for the genus *Cynopterus* based on this study and Campbell et al. (2004), along with private mutations observed in the cytochrome *b* gene.

Comparison of mitochondrial and genome-wide trees highlighted instances of mito-nuclear discordance pointing to past introgression (Fig. 1). For example, one individual of *C. minutus* carried the mitochondrial DNA haplotype of the larger sympatric species *C. princeps* (Fig. 1; Table 1). Similarly, one of the individuals from the undescribed Palawan lineage carried the mtDNA haplotype of *C. brachyotis* (*sensu stricto*) (Fig. 1). Finally, multiple *C. brachyotis* (*sensu stricto*) individuals from the Agartala region in northeastern India and Myanmar (data from Campbell et al. 2004) carried a mitochondrial DNA haplotype that is genetically closer to *C. sphinx* than to *C. brachyotis* (Fig. 1).

As expected, the nuclear data provided better resolution than the mitochondrial cyt *b* gene with regards to relationships among species, especially deeper in the genus. We observed similar results based on both concatenated and species tree methods (Edwards et al. 2007; Edwards 2009; Liu et al. 2010; Stamatakis 2014) (Fig. 1a and S1). The tree topology was not influenced by the inclusion of historic samples that are subject to DNA degradation (Fig. 1a and S2). We identified private mutations for the putative species-level lineages of *Cynopterus* as summarized in Table 1.

### Introgression and coalescent modelling

We observed multiple instances of introgression based on ABBA-BABA analysis (Green et al. 2010) using ANGSD. Concordant results were obtained for both SNP datasets, suggesting the choice of reference genomes did not bias results (Tables S4 and S5). ANGSD results were further confirmed by ABBA-BABA analysis using DSuite (Table S6). Our f-branch analysis (Malinsky et al. 2021) did not identify gene flow in the ancestral lineage, but in pairwise combinations primarily involving two separate lineages: the undescribed Palawan population and *C. brachyotis* (Fig. S3).

To further investigate instances of genetic introgression, we used the coalescent modelling approach implemented in G-PhoCS, which identified multiple secondary gene flow events and two cases in which the divergence times and their confidence intervals overlapped (Fig. 2). The relationship between *C. luzoniensis* and the two separate, undescribed lineages from Palawan and Sulawesi, respectively, could not be conclusively resolved. Similarly, divergence timing between *C. ceylonensis*, *C. horsfieldii*, and the clade leading to *C. sphinx* and *C. minutus* overlapped. Nodal areas of the tree in which divergence events occurred in quick succession were further examined by switching taxa and re-running G-PhoCS. Switching of taxa did not drastically alter the timing of divergence or the overlap in divergence timing among taxa involved (Fig. S4 and S5).

**Figure 2:**
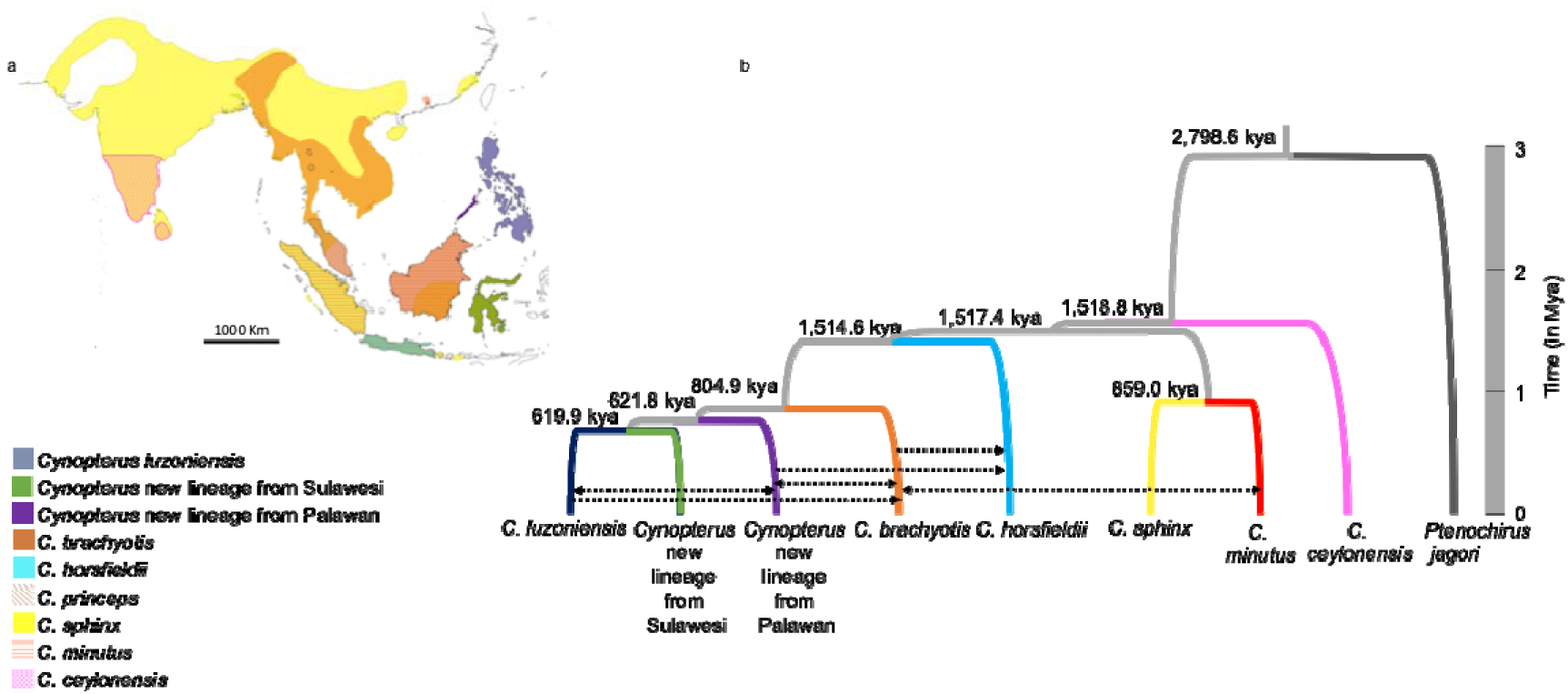
a) Revised distribution of the various lineages of the genus *Cynopterus* highlighting the overlap in range. b) Mean estimates of divergence time and gene flow between various species of the genus *Cynopterus* using the coalescent modelling framework in G-PhoCS (confidence intervals not shown). We analyzed 482 non-linked neutral loci using the topology observed in the RAxML tree (Fig. 1a) for this analysis. Arrows indicate gene flow between species. Only total gene flow over 1% is depicted using black arrows in the figure.

Finally, we used multiple taxon combinations for a comprehensive set of qpbrute analyses. When all taxa including the outgroup (n = 9) were considered, the analysis did not complete even after a year. Therefore, we divided the taxa into two subsets. For the first subset, we selected *C. brachyotis*, *C. sphinx*, *C. minutus*, *C. ceylonensis*, and *Ptenochirus jagori*, another cynopterine fruit bat, as an outgroup. We identified nine possible admixturegraphs in qpBrute, seven of which were equally likely. In the next step, we added *C. horsfieldii* to the seven models and identified 36 possible models, 18 of which were selected after Bayes factor analysis (Kass and Raftery 1995). Two models were discarded due to geographical inconsistencies, leaving 16 plausible models summarized in Figure 3a. For the second subset, we selected *C. luzoniensis*, *C. horsfieldii*, and the two separate undescribed lineages from Palawan and Sulawesi, with *Ptenochirus jagori* as an outgroup. This resulted in 98 models in qpBrute, six of which were selected after Bayes factor analysis, as summarized in Figure 3b.

**Figure 3:**
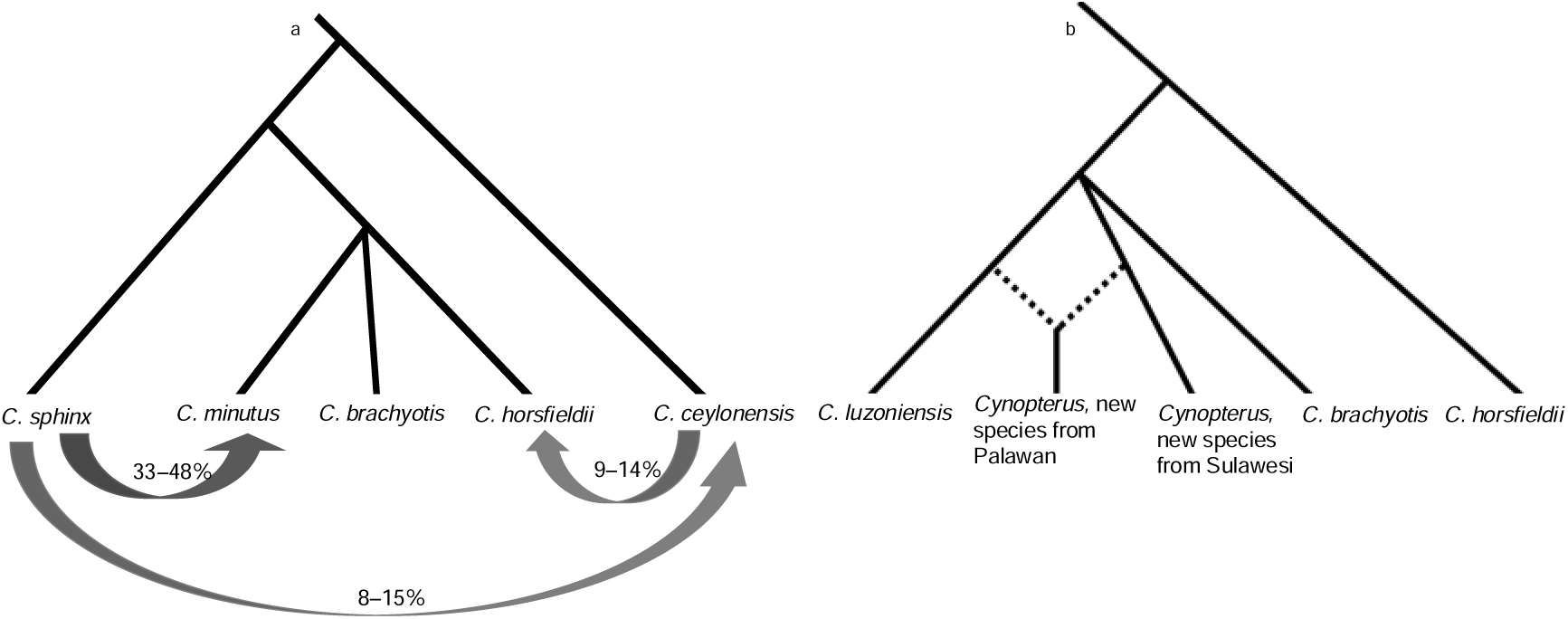
Admixture graph of gene flow dynamics observed in the genus *Cynopterus* using qpBrute. a) Schematic of genetic ancestry and admixture proportions between *C. sphinx*, *C. minutus*, *C. brachyotis*, *C. horsfieldii*, and *C. ceylonensis* based on a combination of 16 best-fit admixture graphs; b) Schematic of genetic ancestry and admixture proportions among *C. luzoniensis*, *C. brachyotis*, *C. horsfieldii* and the two undescribed lineages from Palawan and Sulawesi based on a combination of six best-fit admixture graphs.

### Genes involved in introgression and speciation

A total of 56 species combinations were permissible for sliding window ABBA-BABA analysis based on the RAxML phylogenetic tree. For each comparison, we investigated 7,886 windows spanning 100kb each and considered the subset that contained at least 50 SNPs per window (n = 1180), looking for signatures of introgression. Out of these windows, ∼40% (n = 471) exhibited elevated signatures of either ABBA-like or BABA-like SNPs in at least one of the species combinations tested. Two out of 471 windows were present in at least 50% of the species combinations compared (Fig. 4a). We identified four potential protein coding genes within these windows, one of which belonged to the spermatogenesis-associated protein 46-like family (Fig. 4a).

**Figure 4:**
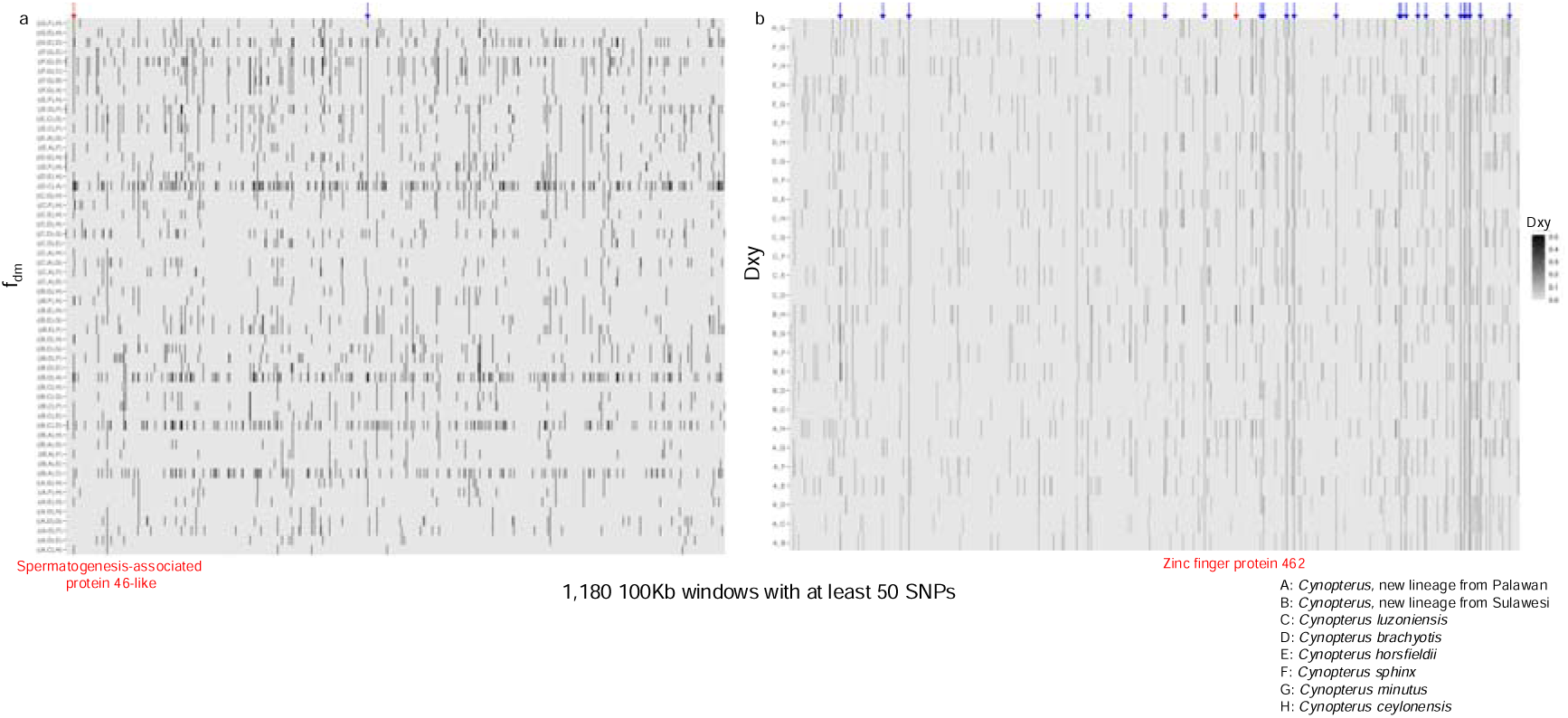
Sliding window analysis across 1,180 suitable windows of 100kb length. a) Only windows with a high introgression signal (top 10%) are colored in black. Two windows with high introgression that are shared among at least 50% of the combinations are marked by arrows. The red arrow depicts the introgression window containing Spermatogenesis-associated protein 46. b) Windows of pairwise divergence between the various lineages of the genus *Cynopterus*. The arrows mark high divergence windows that are shared among at least 50% of the combinations tested. The red arrow marks the window containing the Zinc finger protein 462, which plays a role in cranial morphology.

Similarly, for the analysis of pairwise differentiation (Dxy) across the genome, we identified 28 windows with an elevated Dxy signal present in at least 50% of the combinations compared (Fig. 4b). We identified 16 genes within these windows (Table S7); of these, the Zinc finger protein 462 (*ZNF462*) is known to play an important role in cranial morphology (Weiss et al. 2017; Kruszka et al. 2019). We did not observe significant enrichment of ontology terms (Fig. 4b).

### Ancestry painting

We used ancestry painting to examine the genomic contributions from possible parental species into daughter lineages for two cases of putative hybrid speciation inferred from introgression analysis. We identified 3,120 SNPs that were fixed for different alleles in *C. brachyotis* and *C. sphinx*, 2,879 of which were used for ancestry painting (Fu et al. 2015). In our analysis of the admixed origin of *C. minutus*, we observed a higher allelic contribution of *C. sphinx* when compared to *C. brachyotis* (Fig. 5a). Further, we identified 349 genes associated with fixed SNPs, for which gene ontology analysis identified an enrichment for genes involved in platelet activation and axon guidance, among others (Table 2). In comparison, we observed only 487 SNPs fixed for different alleles between *C. luzoniensis* and *C. brachyotis*, 469 of which were used for analysis into the mixed origin of the undescribed Palawan lineage. Interestingly, the Palawan lineage appears to be a hybrid species with nearly equal contributions from *C. brachyotis* and *C. luzoniensis* (Fig. 5b). We identified 120 genes associated with fixed SNPs but failed to find an enrichment of ontology terms within this subset.

**Figure 5:**
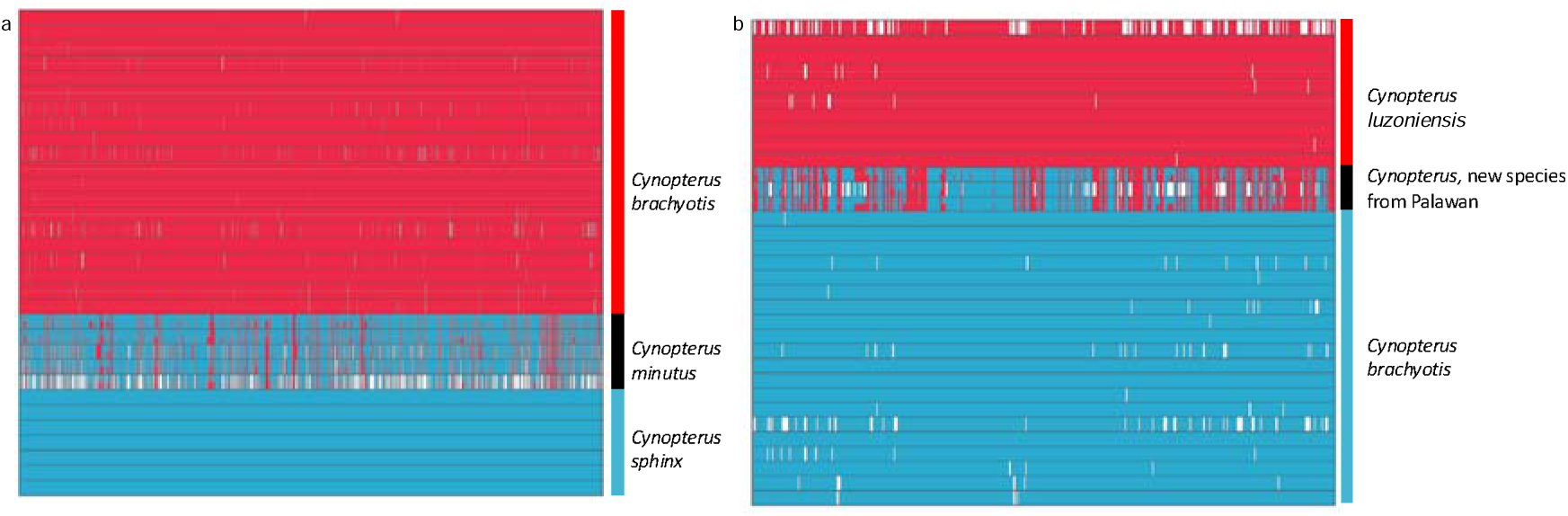
Ancestry painting of putative hybrid species based on loci which are fixed for different alleles in the parental species. a) Parental species: *Cynopterus brachyotis* and *C. sphinx*; putative hybrid species *C. minutus*; b) Parental species: *Cynopterus brachyotis* and *C. luzoniensis*; putative hybrid species is an undescribed lineage from Palawan.

**Table 2:** Gene ontology enrichment analysis of the regions fixed for different alleles in *Cynopterus sphinx* and *C. brachyotis*

| Enrichment False<br>Discovery Rate | Number of<br>Genes | Pathway<br>Genes | Fold Enrichment | Pathway |
| --- | --- | --- | --- | --- |
| 0.0048442247890527 | 8 | 124 | 6.9373097991479 | Platelet<br>activation |
| 0.0342388095183431 | 8 | 181 | 4.75263212759304 | Axon guidance |
| 0.0357639297934434 | 8 | 210 | 4.096316262354 | Rap1 signaling<br>pathway |
| 0.0357639297934434 | 7 | 166 | 4.53432598317799 | CGMP-PKG<br>signaling<br>pathway |
| 0.0357639297934434 | 6 | 120 | 5.37641509433962 | AMPK<br>signaling<br>pathway |
| 0.0357639297934434 | 6 | 113 | 5.70946735682084 | Cholinergic<br>synapse |
| 0.0358450373843572 | 4 | 49 | 8.77782056218714 | Cocaine<br>addiction |
| 0.0478665088471901 | 5 | 92 | 5.84392945036916 | Salivary<br>secretion |

## DISCUSSION

Reproductive and geographical isolation have long been regarded as important requirements for successful speciation (Coyne and Orr 2004). The allopatric speciation paradigm posits that the most widespread mechanism of differentiation is the geographic separation of two daughter lineages, which – in the absence of gene flow – slowly diverge into independent species (Coyne and Orr 2004; Wiens 2004). Gene flow is a powerful homogenizing agent, and classical population genetics dictates that less than a handful of gene flow events per generation suffice to keep two populations homogeneous and panmictic (Hartl and Clark 1997). Consequently, hybridization and introgression have long been viewed as rare, exceptional and/or counterproductive to differentiation. With the advent of Next-Generation Sequencing, we know that introgression is much more widespread in nature than previously thought (Green et al. 2010; Harrison and Larson 2014; Chattopadhyay et al. 2016; Garg et al. 2018; Taylor and Larson 2019; Cros et al. 2020). Even so, it remains unclear whether introgression is merely an incidental side-product of the diversification process, or whether allelic exchange between species can actively contribute to diversification (Green et al. 2010; Rossi et al. 2024).

Our genome-wide dataset of pteropodid fruit bats centered around the *C. brachyotis* species complex unveiled a diverse radiation whose species richness has likely been underestimated. Taxa traditionally assigned to *C. brachyotis* divided into six deep lineages that are arguably at the species level (Fig. 1; Table 1). More importantly, two of them (*C. minutus, C. ceylonensis*) emerged outside the main clade of *C. brachyotis* taxa and in positions more closely related to other traditional *Cynopterus* species. Two of the species-level lineages we identified (one from Palawan and one from Sulawesi) currently lack a scientific name, to the best of our knowledge, and await scientific description and morphological characterization (Fig. 1; Table 1). A full taxonomic revision of the entire genus would be fruitful and may uncover even more species-level diversity.

A simple comparison between phylogenetic trees based on nuclear versus mitochondrial data revealed multiple incongruences that are probably not based on neutral processes such as incomplete lineage sorting, but instead bear the hallmarks of introgressive gene flow between species (Fig. 1). This includes members of *C. minutus, C. brachyotis* and the unnamed Palawan lineage that carry the mitochondrial haplotype of geographically adjacent or overlapping *Cynopterus* species. To test whether genetic introgression may have accounted for such patterns, we carried out a slew of gene flow analyses, ranging from simple ABBA-BABA computations (Table S4-S6) and f-branch statistics (Fig. S3) to coalescent modelling using G-PhoCS (Fig. 2) and admixture graph analysis using qpBrute (Fig. 3). These analyses showed that our radiation of *Cynopterus* bats has undergone rampant introgression during its diversification (Fig. 2, 3b, and 5). Depending on the analysis, fewer or more gene flow events were identified, with the most significant ones consistently involving the following two parts of the tree: (1) The unnamed lineage from Palawan, an island bridging Borneo with the main islands of the Philippines, showed variable gene flow contributions from *C. luzoniensis* (from the main Philippines) and *C. brachyotis* (which occurs on Borneo); and (2) depending on analysis, *C. minutus* emerged as sister of *C. sphinx* or *C. brachyotis*, with introgressive gene flow from the opposite species, respectively (Fig. 1, 2b, 3, and 5).

Chromosome painting further shed light on the potential hybrid speciation origin of *C. minutus* and the Palawan lineage, respectively (Fig. 5), revealing that the Palawan lineage roughly carries a 50:50 mix of allelic contributions from the Philippine *C. luzoniensis* versus Bornean *C. brachyotis*, consistent with the geographic position of Palawan in between Borneo and the northern Philippines (Fig. 5b). Palawan is considered a springboard for diversity in the Philippines allowing for exchange of taxa from the Sunda shelf (Bird et al. 2005; Blackburn et al. 2010; Esselstyn et al. 2010). In contrast, the Sundaic rainforest species *C. minutus* shows a relatively minor influx of alleles from the sympatric *C. brachyotis*, and instead is largely composed of allelic contributions from *C. sphinx*, a species from the subtropical monsoon belt that is not currently known to overlap with *C. minutus* (Fig. 5a). Hence, our analyses indicate that introgressive hybridization may act in different ways to shape *Cynopterus* species diversity: in the case of the Palawan lineage, the bat population evolving in this relatively small landmass has been the product of colonization waves of roughly equal proportions from two nearby landmasses; in the case of *C. minutus*, a widespread Sundaic rainforest lineage may have undergone differentiation from neighboring populations to the north adapted to drier habitats (i.e., *C. sphinx*) via introgression from an overlapping Sundaic bat (Fig. 2, and 5a).

Given the potential significance of functional locus transfers between species in *Cynopterus* diversification, we performed comprehensive genomic scans across all 56 possible pairwise species combinations to flag two types of genomic areas: (1) genomic windows with a high signal of introgression (these are typically windows in which two species exhibit less differentiation than we would expect based on the tree); and (2) genomic windows with extremely high differentiation. Of nearly 7900 windows screened, only 278 (i.e., ∼3.5%) displayed an introgression signal in at least one of the pairwise comparisons. We only considered windows with top 10% signatures of introgression. Two of these windows were shared across at least 50% of combinations. The most commonly occurring window displayed a signature of introgression in all possible combinations except one, with 34 of the combinations exhibiting top 10% fdM values, and refers to a genomic window containing a gene belonging to the spermatogenesis-associated protein 46-like family (*Spata46*) (Fig. 4a). In mice, *Spata46* plays an important role in sperm-egg fusion and sperm head structure. Deficiency of this protein leads to subfertility in mice, supporting its essential role in fertility (Chen et al. 2016). The pronounced signature of introgression in the window containing a gene from this family across virtually all possible taxon combinations suggests that this protein may be implicated in male fertility in *Cynopterus* and may facilitate some of the successful introgression and even hybrid speciation events observed in the genus (Fig. 5). Among the genomic regions exhibiting signatures of divergence between species pairs we identified a window of high divergence containing the Zinc finger protein 462 (*ZNF462*) (Fig. 4b), which plays a critical role in embryogenesis and in determining cranial morphology (Weiss et al. 2017; Kruszka et al. 2019). Skull morphology is a major distinguishing feature in many cryptic bat species, including *Cynopterus* (Bates and Harrison 1997). Given the high level of divergence observed in and around this gene, it may play an important role in *Cynopterus* species divergence and differentiation.

In summary, our analyses suggest that *Cynopterus* fruit bats form a cryptic radiation of species which continue to be subject to occasional gene flow among each other. We show that – far from being an impediment to diversification or counteracting the speciation process – introgressive gene flow has likely boosted the diversification potential of the genus. We detected two possible hybrid species with variable allelic contribution percentages from their putative parental species, and uncovered introgressive gene flow events in several more. Our gene scans identified loci implicated in fertility that may be involved in facilitating genetic introgression and possibly even hybrid speciation. More thorough genomic sampling across many additional organismic radiations in the future may corroborate that the patterns we unveiled in *Cynopterus* are a more general feature of the speciation process.

## AVAILABILITY OF DATA AND MATERIALS

All the original data necessary to reproduce the analyses reported in this study can be accessed through the Dryad link: http://datadryad.org/share/14Juhqaeoj6RwM5o13ZoAzdhotpZb8N_9ld4614yRhU

The raw sequence data generated as part of this study is available on NCBI under Bioproject PRJNA1178127.

## ACKNOWLEDGMENTS

We thank Lawrence Heaney at the Field Museum, Chicago, USA; Marcus Chua and Kelvin Lim at the Lee Kong Chian Natural History Museum (LKCNHM), Singapore; Museum Zoologicum Bogoriense, Cibinong, West Java, Indonesia (MZB); Carlos Urdiales at the Estación Biológica de Doñana (EBD) for providing samples for this study. BC acknowledges the support provided by Uma Ramakrishnan and Pilot Dovih for helping with sample processing. BC acknowledges the support provided by Giovanni Forcina in procuring samples from EBD. Permissions for research, fieldwork, and collection were granted by the Indonesian Ministry of Environment and Forestry, RISTEK-BRIN (formerly RISTEK) and local government entities (1337/FRP/SM/IV/2012, 387/SIP/FRP/SM/IX/2012, 396/FRP/SM/II/2014). SMT and SW facilitated collaborative fieldwork and research through a Memorandum of Understanding between the City College of New York (CCNY) and the Museum Zoologicum Bogoriense (MZB). We are indebted to Abdul Rahman, the late Edy Toyibi, Sheherazade, Yusep Synata, and the staff of BKSDA and MZB for their assistance. KMG acknowledges the help provided by Evan K. Irving-Pease for running qpBrute.

## AUTHOR CONTRIBUTIONS

B.C., K.M.G., and F.E.R.: Conceptualization; B.C. and K.M.G.: Formal analysis; S.M.T., J.L.C., R.P.J., F.A.B.A.K., S.W., I.H.M., and G.J.D.S.: Resources; and B.C., K.M.G., and F.E.R.: Writing– original draft preparation; all authors: Writing–review and editing.

## SUPPORTING INFORMATION

Supporting tables S1–S7

Supporting figures S1–S5

