## Supporting information for "Genetic introgression is a catalyst for diversification in an Old World fruit bat radiation"

**SUPPORTING TABLES**

Table S1: Details of the samples used in this study. Abbreviations: LKCNHM: Lee Kong Chian Natural History Museum, Singapore; MZB: Museum Zoologicum Bogoriense, Cibinong, West Java, Indonesia; NA: not applicable (Excel sheet).

Table S2: Details of the sequences obtained from GenBank included for cytochrome *b* phylogenetic analysis (Excel sheet)

Table S3: Summary of the number of SNPs isolated and filtered for the two SNP datasets.

| Dataset | Number of unfiltered SNPs | Number of SNPs removed due to linkage | Number of SNPs removed that were not in Hardy-Weinberg equilibrium | Number of non-neutral SNPs removed | Number of SNPs retained | Overall level of missing data |
| --- | --- | --- | --- | --- | --- | --- |
| SNP dataset I: SNPs obtained after mapping to *Cynopterus brachyotis* scaffold level assembly | 331,534 | 89,188 | 6,720 | 18,274 | 217,352 | 6.6% |
| SNP dataset II: SNPs obtained after mapping to pseudo-chromosomal assembly of *C. brachyotis* | 172,049 | 50,093 | 4,406 | 9,714 | 107,836 | 6.5% |

Table S4: ABBA-BABA results based on SNP dataset I estimated using ANGSD, where H1, H2, H3 and H4 are the four species compared for ABBA-BABA analysis; D – Patterson’s D statistic; Z – Z score; nABBA – number of ABBA patterns observed; nBABA – number of BABA patterns observed; nBlocks – number of blocks with observed data. (Excel sheet)

Table S5: ABBA-BABA results based on SNP dataset II estimated using ANGSD, where H1, H2, H3 and H4 are the four species compared for ABBA-BABA analysis; D – Patterson’s D statistic; Z – Z score; nABBA – number of ABBA patterns observed; nBABA – number of BABA patterns observed; nBlocks – number of blocks with observed data. (Excel sheet)

Table S6: ABBA-BABA results based on SNP dataset I estimated using DSuite, where P1, P2, and P3 are the species compared for ABBA-BABA analysis; Dstatistic– Patterson’s D statistic; BBAA – number of BBAA patterns observed; ABBA – number of ABBA patterns observed; BABA – number of BABA patterns observed. (Excel sheet)

Table S7: List of genes identified in 28 windows of high divergence (Dxy) which are present in at least 50% of the species combinations investigated in the genus *Cynopterus*.

| Gene ID | Annotated function | Gene symbol |
| --- | --- | --- |
| ID=g6538 | Ganglioside-induced differentiation-associated protein 1 | GDAP1 |
| ID=g12202 | NA |  |
| ID=g12201 | Roundabout | ROBO4 |
| ID=g12909 | NA |  |
| ID=g14458 | NA |  |
| ID=g16925 | NA |  |
| ID=g18948 | cytochrome P450 | P450 |
| ID=g19582 | NA |  |
| ID=g2679 | NA |  |
| ID=g2678 | Otoferlin | OTOF |
| ID=g2680 | Dynein regulatory complex | DRC1 |
| ID=g8846 | Cilia and flagella associated protein 61 | CFAP61 |
| ID=g8847 | Ral GTPase-activating protein subunit alpha-2 | RALGAPA2 |
| ID=g11400 | Zinc finger protein 462 | ZNF462 |
| ID=g16305 | Signal transducer and activator of transcription | STAT6 |
| ID=g16306 | lipoprotein receptor-related protein 1 | LRP1 |

**SUPPORTING FIGURES**

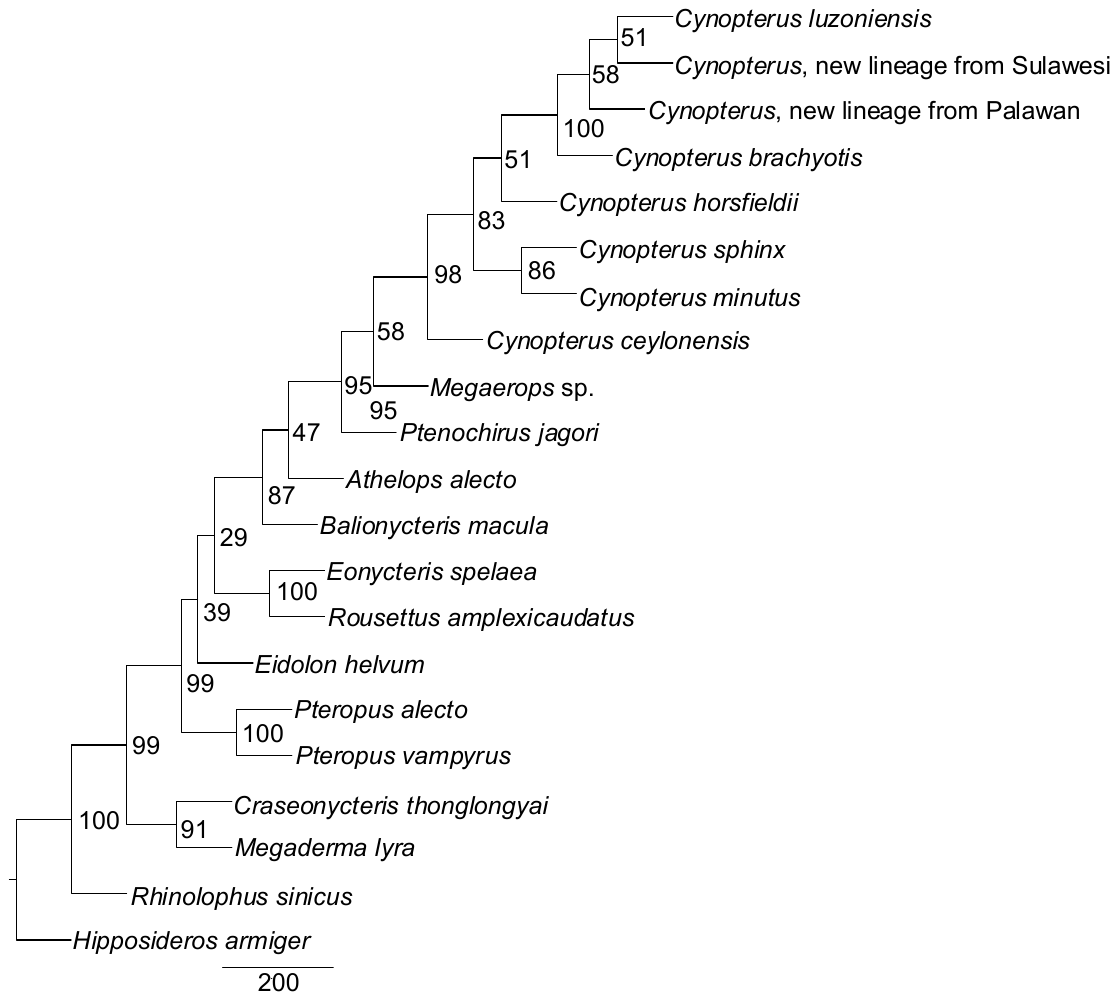

Figure S1: Cladogram of various members of the genus *Cynopterus* based on 1,182 nuclear loci. MP-EST was used to construct this species tree.

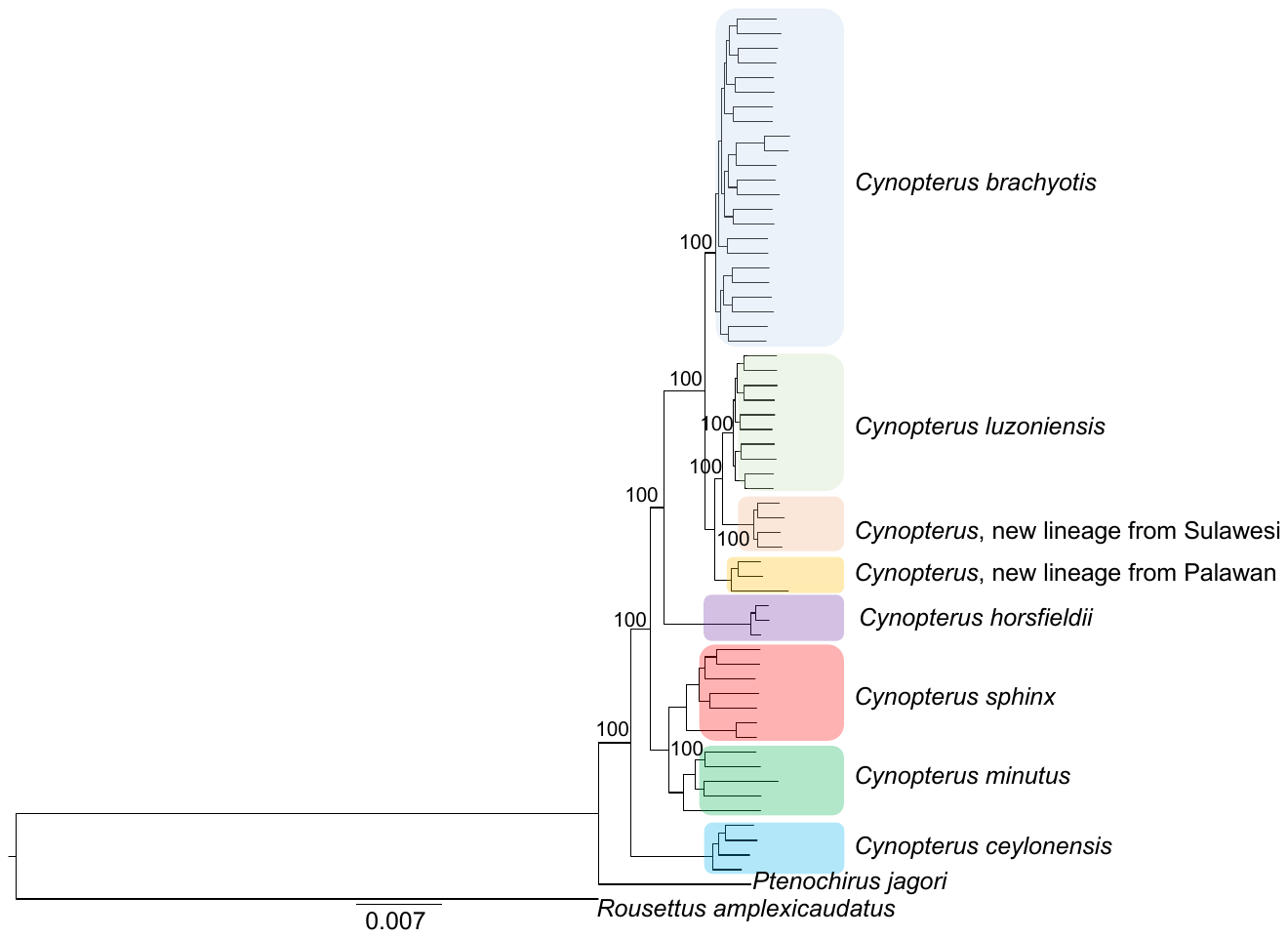

Figure S2: Maximum likelihood tree based on concatenated nuclear loci (number of loci: 1,182; total length of the sequence 1,466,886 bp) generated using RAxML. Only fresh samples were used for the phylogenetic reconstruction. Nodal values indicate bootstrap support. Support values are shown only for major nodes.

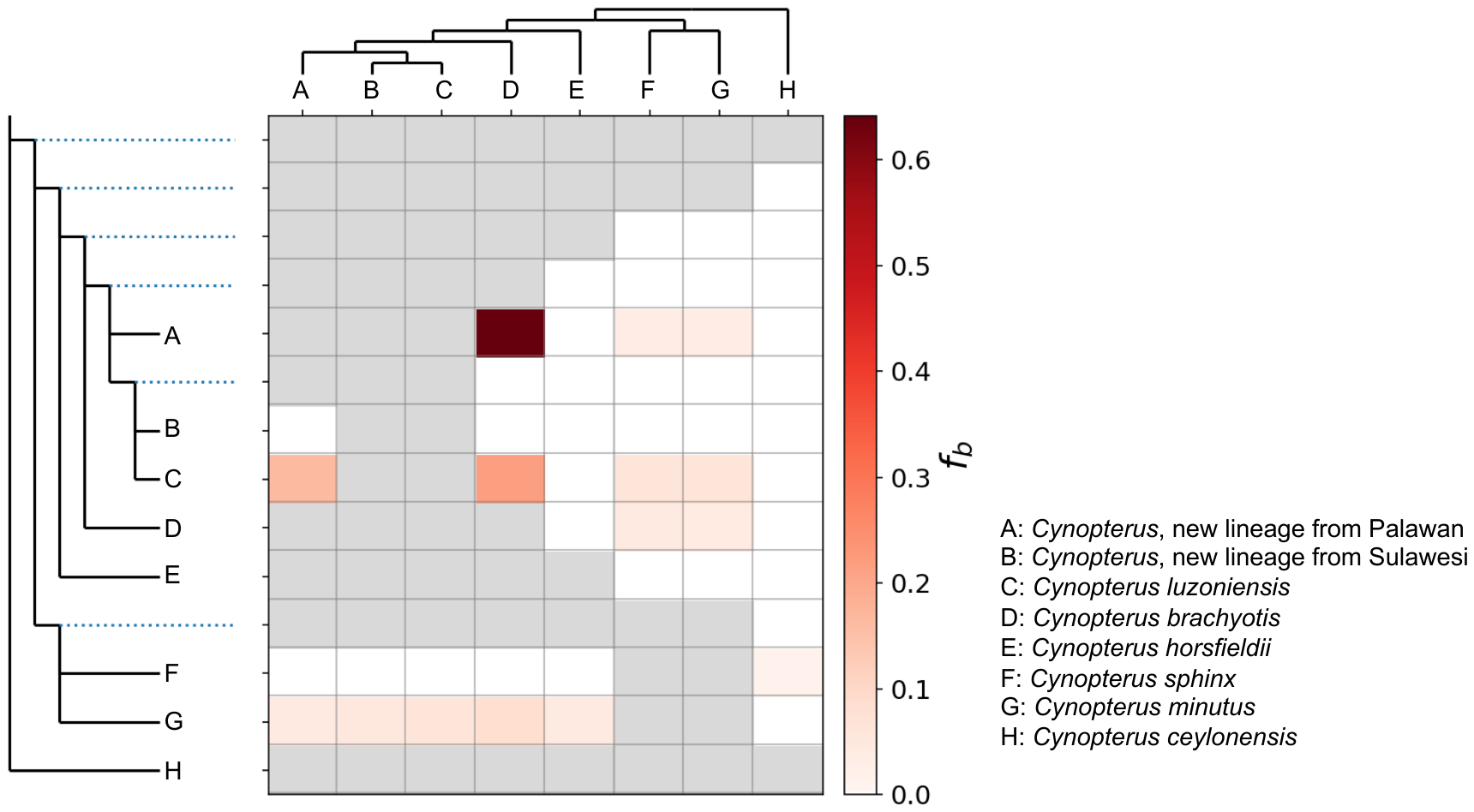

Figure S3: f-branch statistics indicating introgression between various lineages of the genus *Cynopterus*.

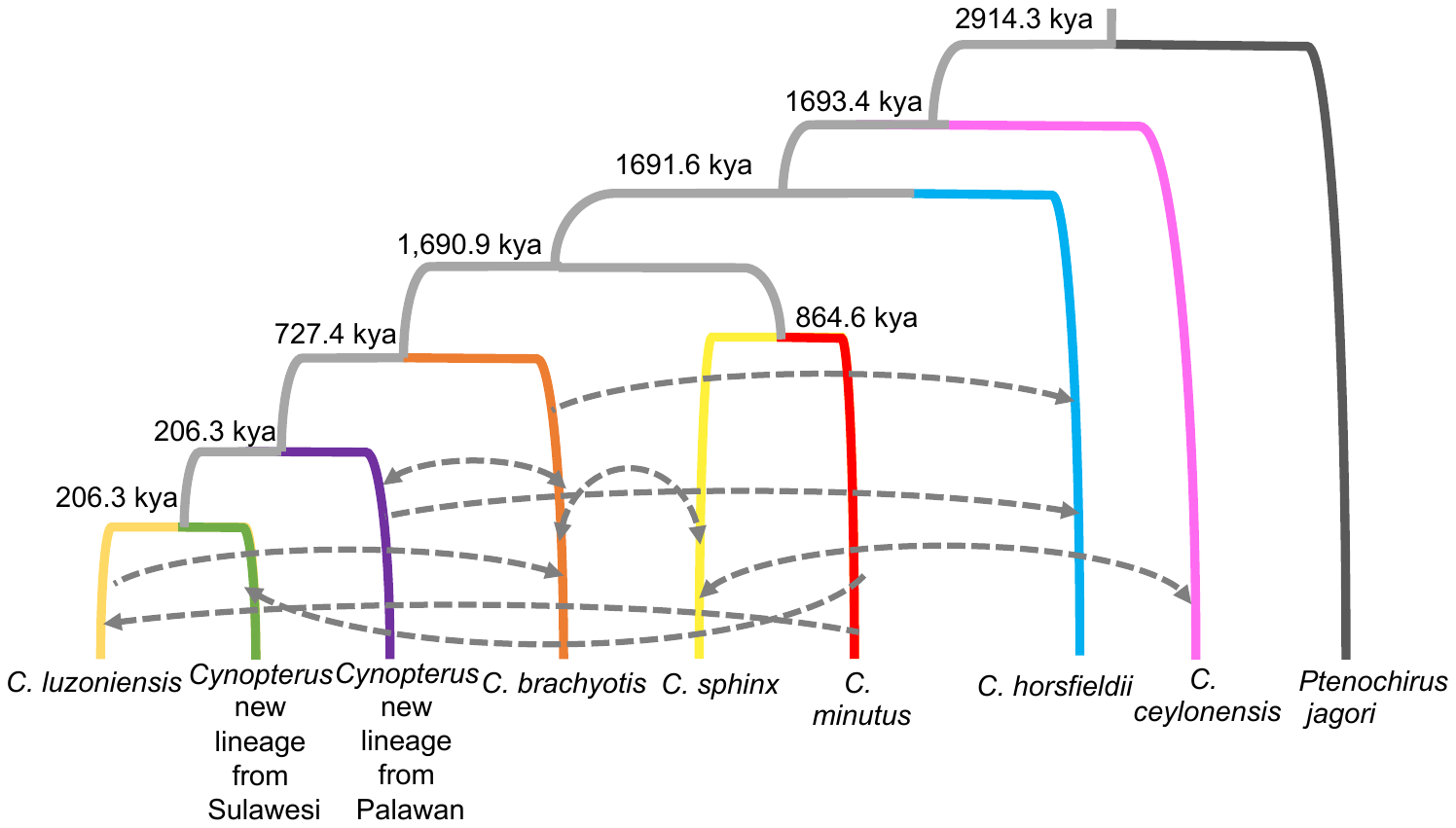

Figure S4: Estimates of divergence time and gene flow between various species of the genus *Cynopterus* using the coalescent modelling framework in G-PhoCS. We changed the topology (switching the relationship between *C. sphinx*, *C. minutus*, and *C. horsfieldii*) for this analysis, using 482 non-linked neutral loci. Arrows indicate gene flow between species. Grey arrows indicate that gene flow results are stable across runs.

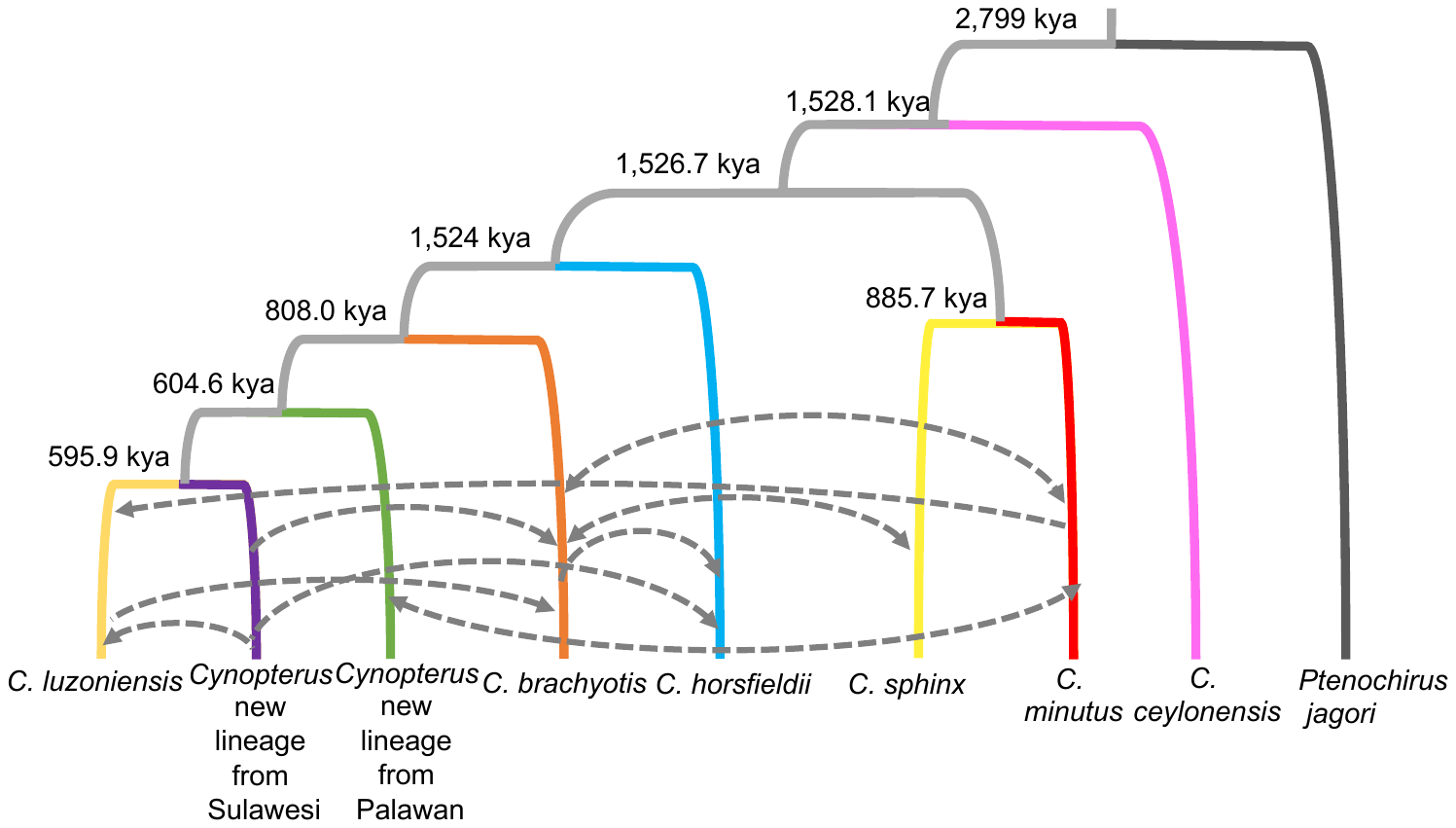

Figure S5: Estimates of divergence time and gene flow between various species of the genus *Cynopterus* using the coalescent modelling framework in G-PhoCS. We changed the topology (switching the relationship between *C. luzoniensis* and the two undescribed lineages from Palawan and Sulawesi, respectively) for this analysis, using 482 non-linked neutral loci. Arrows indicate gene flow between species. Grey arrows indicate that gene flow results are stable across runs.
